# Distinct modes of heat shock transcription factor interactions with mitotic chromosomes

**DOI:** 10.1101/2022.10.05.511012

**Authors:** Rachel M. Price, Marek A. Budzyński, Junzhou Shen, Jennifer E. Mitchell, James Z.J. Kwan, Sheila S. Teves

## Abstract

A large number of transcription factors have been shown to bind and interact with mitotic chromosomes, which may promote the efficient reactivation of transcriptional programs following cell division. Although the DNA-binding domain (DBD) contributes strongly to TF behavior, TFs from the same DBD family can display distinct binding behaviors during mitosis. To define the mechanisms governing TF behavior during mitosis in mouse embryonic stem cells, we examined two related TFs: Heat Shock Factor 1 and 2 (HSF1 and HSF2). We found that HSF2 maintains site-specific binding genome-wide during mitosis, whereas HSF1 binding is globally decreased. Surprisingly, live-cell imaging shows that both factors appear excluded from mitotic chromosomes, and are similarly more dynamic in mitosis than in interphase. Exclusion from mitotic DNA is not due to extrinsic factors like nuclear import and export mechanisms. Rather, we found that the HSF2 DBD alone can coat mitotic chromosomes, but is insufficient to promote HSF1 coating. These data further confirm that site-specific binding and chromosome coating are independent properties, and that for some TFs, mitotic behavior is largely determined by the non-DBD regions.

## Introduction

Mitosis is accompanied by major changes in cellular morphology and function. At the onset of M phase, DNA is globally condensed and reorganized to form characteristic mitotic chromosomes (1), and transcription from mitotic chromosomes is largely shut down (2). The nuclear membrane also loses its continuity in mitotic cells and no longer serves as a tight barrier between the cytoplasm and chromatin (3). Given that transcription is rapidly re-established after cell division (2, 4), mechanisms must exist to facilitate the precise and accurate reestablishment of gene expression after mitosis (5). One proposed mechanism, termed mitotic bookmarking, suggests that the binding of proteins, mainly transcription factors (TFs), to specific genomic loci during mitosis marks genes for rapid activation after cell division (6).

Early studies using chromatin immunoprecipitation (ChIP)-based methods identified only a few TFs that bind to specific loci during mitosis, including HSF2, TFIID, and TFIIB (7, 8). Another study using immunofluorescence reported that most TFs are displaced from mitotic chromatin, further supporting the hypothesis that mitotic bookmarking is rare (9). This rarity was due in part to formaldehyde fixation, which was found to artificially evict proteins from DNA during mitosis (10). Indeed, a recent study using ChIP-based methods with alternative fixatives detected more site-specific binding during mitosis by various TFs (11), and advances in live-cell imaging led to the identification of an increasing number of TFs that are enriched on mitotic DNA, including GATA1, FOXA1, SOX2, TBP, CTCF, KLF4, and ESRRB (10, 12–17). Since then, studies using imaging and proteomics approaches have identified hundreds of proteins, including TFs, that interact with DNA during mitosis (18–20). A large screen of TF localization during mitosis by live-cell imaging reported varying degrees of DNA-TF interactions, from strongly enriched on mitotic DNA, through moderately enriched, to clearly excluded (18). These studies show that mitotic DNA-TF interactions are best measured by live-cell imaging and/or by chromatin-binding assays that avoid formaldehyde fixation, with the assumption that the chromatin association measured by one method corresponds with the association measured by the other. However, chromatin-binding assays measure site-specific binding of TFs during mitosis, whereas live-cell imaging techniques visualize global mitotic chromatin ‘coating’, which is heavily influenced by non-specific interactions between the TF and the DNA (11, 15). Indeed, some proteins that are unable to bind DNA site-specifically still demonstrate global mitotic coating as visualized by live imaging (10, 11, 15). Although coating is likely a function of both site-specific binding and non-specific interactions (10, 15), live-cell imaging cannot provide information about TF binding at specific loci.

Regardless of the method of measurement, certain TF domains are critical for DNA-TF interaction during mitosis. The DNA-binding domain (DBD) has been shown to drive much of TF behavior (21–23), and would be expected to play a major role in mitotic DNA-TF interactions as well. Indeed, previous work by Raccaud et al suggests that the type of DBD predicts the strength of DNA-TF interactions to a certain degree (18). Additionally, the DBD is necessary for mitotic bookmarking in some cases (10). Yet, TFs with highly similar DBDs often interact differently with mitotic DNA (18, 24), and the mechanisms that drive TF interactions with mitotic DNA remain largely unknown. Even less understood than the DBD is the role of the nuclear import mechanism and the nuclear localization signal (NLS) in mitotic DNA-TF interactions (10, 18, 25). Both the NLS and active nuclear import have been suggested to promote mitotic DNA-protein association. Mutating the NLS of SOX2 resulted in eviction of the TF from mitotic DNA in mESCs, while adding the simian virus 40 (SV40) NLS–a strong viral nuclear localization signal (26)–to HaloTag alone promoted coating of the mitotic DNA (10). However, other work suggested that addition of positively charged amino acids was sufficient to localize proteins to mitotic chromosomes, with or without active nuclear import (18). Additionally, importin-β was shown to be involved in the retention of HNF1β on mitotic DNA in MDCK cells, indicating a link between active nuclear import and mitotic DNA-TF interactions (25).

An example of a pair of TFs with highly similar DBDs but distinct mitotic behaviors is Heat Shock Factors 1 and 2 (HSF1 and HSF2), TFs that belong to the same family based on their largely conserved DBDs (Fig. S1A) (27). Known as the primary regulators of the evolutionarily conserved Heat Shock Response defense mechanism, activated HSFs trimerize and drive the expression of genes in response to protein-damaging stress (28). In vertebrates, HSF1 and HSF2 are ubiquitously expressed in all tissues (29), and have been shown to form homo- and hetero-trimers upon activation (30). HSF2 was one of the earliest TFs identified to bind to mitotic DNA at specific loci (7, 24). In contrast, eviction of HSF1 from mitotic chromatin was established by multiple studies including those that used live-cell imaging approaches (9, 10, 24).

Using a combination of genomics and live-cell imaging in mESCs, we investigate the molecular underpinnings behind the interactions of HSFs with mitotic DNA. We found that, although endogenous HSF2 binds site-specifically to mitotic DNA via Cleavage Under Targets and Tagmentation (CUT&Tag) (31) whereas HSF1 does not, live-cell imaging shows that both HaloTagged HSF1 and HSF2 do not associate with mitotic chromatin. Despite their differences in site-specific binding, the dynamics of HSF1-Halo and Halo-HSF2 are remarkably similar when examined with single-molecule live-cell imaging and single-particle tracking (SPT). Additionally, we find that the SV40 NLS is unable to induce mitotic coating of Halo-HSF2, though it does so for HaloTag alone. However, the truncated HSF2 DBD is able to coat mitotic DNA, and this coating is inhibited by the heptad repeats A and B (HR A/B) within HSF2. Though able to coat in isolation, the HSF2 DBD is unable to induce coating of the non-DBD domains of HSF1. Therefore, we demonstrate that the non-DBD regions of HSF1 and HSF2 dictate their exclusion from mitotic DNA, in contrast to other TFs such as SOX2 and SOX13 whose enrichment is driven almost entirely by the DBD.

## Results

### HSF1 and HSF2 display different modes of site-specific DNA interaction during mitosis

To investigate the binding behaviors of endogenous HSF1 and HSF2 in mESCs during mitosis without crosslinking, we performed spike-in normalized CUT&Tag, a recently developed assay that maps protein binding to DNA in native conditions (31). We collected synchronized mitotic cells by treating mESCs with nocodazole, a tubulin polymerase inhibitor (47), for 6 hours followed by shake-off. We then assessed the purity of mitotic cells first by propidium iodine staining followed by flow cytometry, and second by counting DAPI-stained cells. In both assays the mitotic index (percentage of mitotic cells) was above 91% (Fig. S1B-D).

First, we examined SOX2, a transcription factor with established mitotic DNA-binding activity (10, 11, 13, 48), although with some discrepancies between studies. CUT&Tag analysis shows high enrichment of SOX2 at known binding sites, such as at the proximal enhancer of the *Pou5f1* gene in asynchronous samples, and this high level of binding is maintained in mitotic cells (Fig. 1A) with high reproducibility between replicates (Fig. S2). To assess mitotic SOX2 binding genome-wide, we plotted the CUT&Tag signal for all known SOX2 binding sites as heatmaps (Fig. 1B), and displayed the normalized read counts per binding site for asynchronous and mitotic cells as a scatter plot (Fig. 1C). Both heatmap and scatter plot analyses show that SOX2 maintains a high level of site-specific binding in mitosis genome-wide, consistent with previously published studies (13). Therefore, CUT&Tag can measure endogenous TF binding during mitosis.

**Figure 1:**
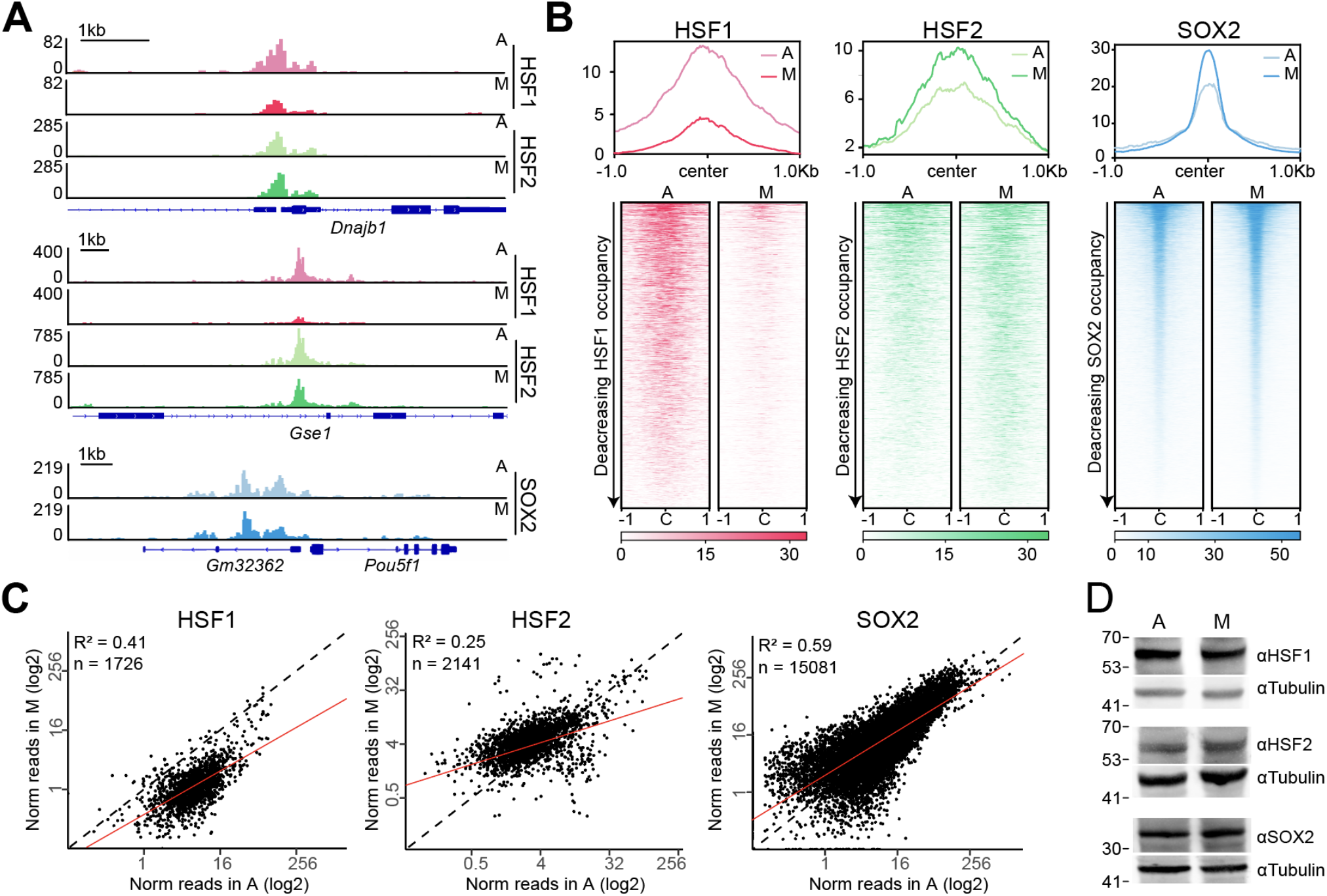
HSF1 and HSF2 have distinct DNA-binding activity in mitosis. **(A)** Gene browser tracks of HSF1 and HSF2 over *Dnajb1* promoter (top) and *Gse1* loci (middle), and SOX2 over *Pou5f1* (OCT4) promoter (bottom) in asynchronous (A) and mitotic (M) cells. **(B)** Genome-wide average plots (top) and heatmaps (bottom) of HSF1 (left), HSF2 (middle), and SOX2 (right) CUT&Tag in asynchronous (A) and mitotic (M) cells. CUT&Tag signal was calculated in a 2 kb window surrounding binding sites of the respective TF. For heatmaps, binding sites were ordered by decreasing occupancy of the given TF. **(C)** Normalized read counts of HSF1 (left), HSF2 (middle), and SOX2 (right) for each binding site in asynchronous (A) and mitotic (M) cells are displayed as a scatter plot. N denotes number of analyzed sites. **(D)** HSF1 (top), HSF2 (middle), and SOX2 (bottom) protein levels in asynchronous (A) and mitotic cells (M). Tubulin is shown as a loading control.

Next, we performed CUT&Tag on HSF1and HSF2 in mitotic and asynchronous cells. We observed strong binding profiles at known HSF1 and HSF2 binding sites in asynchronous cells, including *Dnajb1* and *Gse1* (Fig. 1A). In mitotic samples, we observed decreased HSF1 binding at specific loci, including *Gse1*, whereas HSF2 binding was generally maintained, or even increased, consistent with previous studies (7, 9, 24). To assess HSF1 and HSF2 binding profiles genome-wide, we used SEACR, a CUT&Tag-specific peak calling algorithm, and identified combined binding sites in asynchronous and mitotic samples for HSF1 (1726) and HSF2 (2141) (Fig. S2D). We then plotted HSF1and HSF2 CUT&Tag signal for asynchronous and mitotic samples in a 2-kb region surrounding these identified binding sites as heatmaps and average profiles (Fig. 1A,B). Globally, the average signal of HSF1 binding in mitosis was 6 times lower compared to asynchronous cells (Fig. 1B and S2D). In contrast, HSF2 CUT&Tag signal was maintained or even increased genome-wide during mitosis (Fig. 1B and S2D). To quantify the change in asynchronous versus mitotic signal, we displayed the normalized read counts per binding site as a scatter plot (Fig. 1C). Similar to SOX2, most of the HSF2 signal in mitosis is maintained at similar or higher levels compared to asynchronous population. In contrast, HSF1 signal is largely below the diagonal, showing that for the vast majority of sites, HSF1 binding during mitosis is decreased. The contrasting binding behavior between HSF1 and HSF2 during mitosis is not due to altered protein levels as HSF1, HSF2, and SOX2 show no change in expression during mitosis (Fig. 1D). Our results highlight different modes of site-specific DNA binding for HSF1 and HSF2 during mitosis, despite having highly similar and conserved DBDs.

### HaloTagged HSFs are excluded from mitotic chromosomes when examined with live-cell imaging

Given the differences we observed between mitotic sitespecific binding of HSF1 and HSF2 genome-wide, we next explored how these TFs interact with mitotic chromatin using live-cell imaging. To examine the mitotic coating of HSF1 and HSF2, we stably integrated HaloTagged constructs of each TF expressed under the EF1α promoter (HSF1-Halo, Halo-HSF2, Halo-SOX2; Fig. 2A) into mESCs stably expressing Histone 2B tagged with GFP (H2B-GFP). C-terminal tagging of HSF2 resulted in degradation of the protein, thus it was tagged N-terminally (Fig. S3A). A cell line stably overexpressing HaloTagged SOX2 was used as control for coating behavior. Expression of HaloTagged TF constructs was verified with Western blots against HSF1, HSF2, and SOX2 in asynchronous and mitotic cells (Fig. 2B). HaloTagged TF overexpression cell lines were stained with JF549, a dye which covalently binds to HaloTag (49), and imaged under live-cell conditions. Chromatin enrichment was quantified by calculating the log_2_ ratio of the mean HaloTag signal intensity on mitotic chromosomes over the signal intensity across the whole cell (Fig. 2D). As previously observed, Halo-SOX2 is strongly enriched on mitotic DNA (Fig. 2C,E) in alignment with the high level of specific mitotic binding of endogenous SOX2 genome-wide. Given the reduced specific mitotic binding of endogenous HSF1, we expected and observed that HSF1-Halo is excluded from mitotic DNA (Fig. 2C,E). Surprisingly, Halo-HSF2 is excluded to a similar degree as HSF1-Halo (mean enrichment −0.484 vs −0.458, respectively) (Fig. 2C,E). Exclusion of the HSFs is likely not due to the addition of the HaloTag, which was used to examine the mitotic enrichment of many other TFs in a previous study, including SOX2, SP1, OCT4, and ESRRB (10). In this same study, tagging the aforementioned TFs with an alternative fluorescent tag (mCherry) did not change the chromatin enrichment results. Exclusion of Halo-HSF2 and HSF1-Halo is also not due to fewer putative binding sites for the HSFs, which may influence both specific- and non-specific DNA interactions. A computational prediction of the number of potential binding sites genome-wide using consensus binding motifs revealed roughly similar numbers of putative sites between HSF1 and HSF2 (61,078 and 74,017, respectively), which are also similar to SOX2 (60,812) (Fig. S2E). Therefore, unlike Halo-SOX2, Halo-HSF2 appears excluded from mitotic chromosomes despite high levels of specific mitotic binding by endogenous HSF2 as measured by CUT&Tag (Fig. 1). These results confirm that sitespecific mitotic binding and coating are independent properties that are measurable by different assays (11), and that, at least for HSF2, are driven by distinct mechanisms.

**Figure 2:**
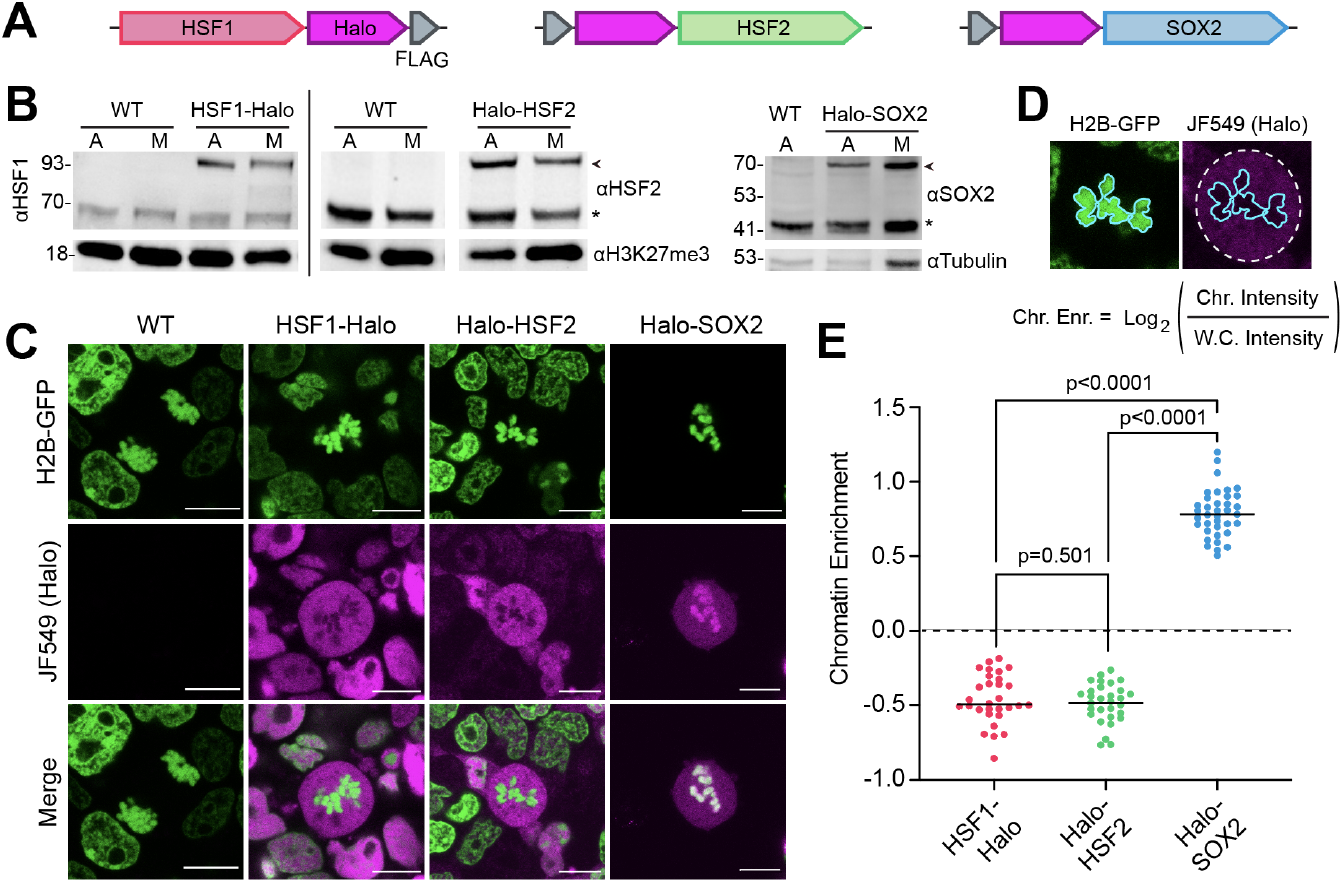
HaloTagged HSFs are excluded from mitotic chromosomes when viewed with live-cell imaging. **(A)** Schematics of stably integrated HaloTag-TF overexpression constructs in WT JM8 cells expressing H2B-GFP. Each TF–SOX2 in blue, HSF1 in red, and HSF2 in green–are tagged with HaloTag (magenta) and FLAG Tag (gray), and expressed under an EF1α promoter. **(B)** Western blots of each HaloTag-TF construct in asynchronous (A) and mitotic (M) cells using α-HSF1 (left), α-HSF2 (middle), and α-SOX2 (right). Mitotic cell populations were synchronized with 50 ng/mL nocodazole for 6 hours before collecting cells via shake-off. Asterisks indicate endogenous TFs (HSF1, HSF2, SOX2). Arrows indicate HaloTagged TFs (Halo-SOX2, HSF1-Halo, and Halo-HSF2). H3K27me3 and Tubulin are shown as loading controls. **(C)** Live-cell fluorescent imaging of HaloTag-TF constructs (magenta) labeled with 200 nM JF549 dye. DNA is visualized with H2B-GFP overexpression (green). Scale bars represent 10 μm. **(D)** Strategy for quantifying TF chromatin enrichment. **(E)** Chromatin enrichment quantification for the indicated HaloTagged TFs (n=30 cells for HSF1-Halo, 30 for Halo-HSF2, and 37 for Halo-Sox2 across 3 biological replicates). Data are visualized as individual data points with mean value indicated. P-value calculated using a standard two-tailed t-test with a 95% confidence interval.

### HaloTagged HSFs are more dynamic in mitosis than in interphase

An alternative method for quantifying DNA-TF interactions is single molecule live-cell imaging coupled with single particle tracking (SPT) (50, 51). Using sparse labeling combined with long exposure times (200 ms; slow-tracking mode), diffusing molecules ‘blur’ out while DNA-interacting molecules appear as diffraction-limited spots (10, 50). Using this method, previous studies have shown that the residence time of Halo-SOX2 molecules on mitotic chromosomes is half of what it is on interphase chromatin (10), suggesting that the interaction of Halo-SOX2 with mitotic DNA is more dynamic despite its ability to coat. To characterize the interaction dynamics of the HSFs with mitotic DNA, we performed SPT on HSF1-Halo in HSF1 knockout (KO) mESCs (HSF1^--/-^; Fig. S3B,C), and on Halo-HSF2 in HSF2 KO mESCs (HSF2^-/-^; Fig. S3D,E) in interphase and mitotic cells. The KO background ensures that the dynamics of the tagged HSFs are independent of the trimerization with untagged endogenous HSFs (52). After image collection, stable molecules are localized and tracked using the SLIMfast algorithm (34). The results are plotted as a log histogram of dwell times as previously described (10, 50) and averaged across three biological replicates (Fig. 3A), with high consistency between replicates (Fig. S4A,B). Cells stably expressing H2B-Halo are used as a photobleaching control as previously described (10) and analyzed similarly. A two-component exponential decay model is fitted to the dwell time curves, from which the time to reach 1% of molecules still bound was calculated (Fig. 3B,C). Both HaloTagged HSFs are more dynamic in mitosis than in interphase, and the dynamics between the two are nearly indistinguishable. The separation between the interphase and mitotic dwell curves is similar, and the time to 1% bound for both HSFs is similarly decreased in mitosis versus interphase (Fig. 3A,B).

**Figure 3:**
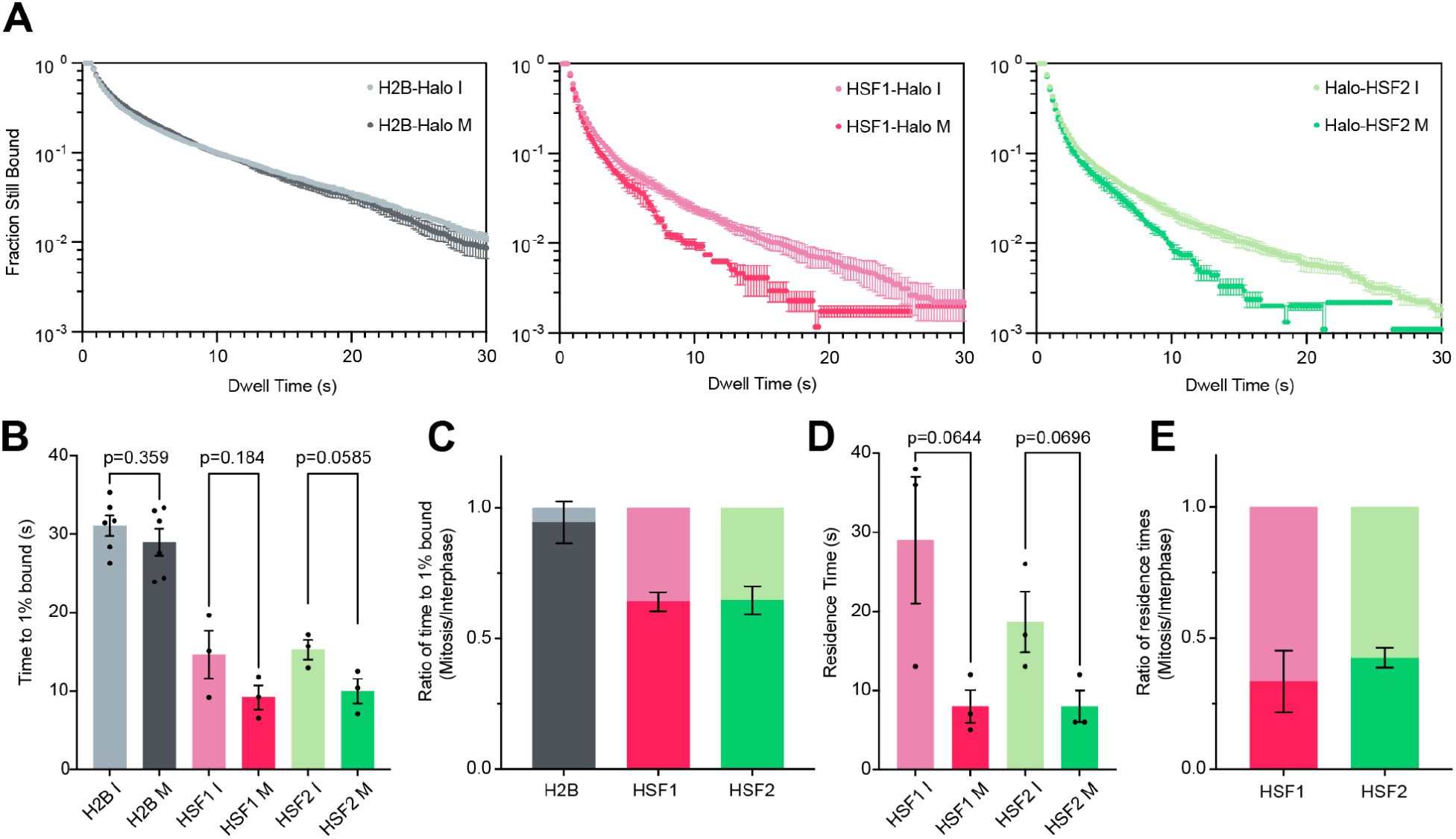
Single-particle tracking reveals that HaloTagged HSF constructs are more dynamic in mitosis than in interphase. **(A)** Dwell time curves of H2B-Halo control (gray, left), HSF1-Halo (red, middle), and Halo-HSF2 (green, right) showing the fraction of HaloTagged molecules bound to DNA in interphase (I, lighter color) and mitosis (M, darker color) over time. Data depicted as mean ±SEM (n=62 cells for H2B-Halo I, 60 for H2B-Halo M, 34 for HSF1-Halo I, 23 for HSF1-Halo M, 32 for Halo-HSF2 I, and 29 for Halo-HSF2 M across three biological replicates). **(B)** Time to 1% of total HaloTagged molecules bound to DNA for each cell line in (A) in interphase and mitosis, where each dot represents one biological replicate. Data depicted as mean ± SEM. **(C)** Ratio of the time to 1% of total HaloTagged molecules bound during mitosis relative to interphase. Data depicted as mean ±SEM. **(D)** Residence times of HSF1-Halo and Halo-HSF2 on DNA in interphase and mitosis, where each dot represents one biological replicate. Data depicted as mean ±SEM. **(E)** Ratio of mitotic residence times for HSF1-Halo and Halo-HSF2 relative to interphase. Data depicted as mean ±SEM. All p-values are calculated using a standard two-tailed t-test with a 95% confidence interval.

To estimate an average residence time for the HaloTagged HSFs, the apparent *k*_off_ for each factor was extracted from the two-component exponential decay model as previously reported (10, 50) and corrected for photobleaching using H2B-Halo (Fig. S4C). The residence time was calculated by taking the inverse of this corrected *k*_off_ (Fig. 3D). Though their overall dynamics are similar, a moderate difference in the residence time in mitosis relative to interphase is observed between HSF1-Halo and Halo-HSF2 (33.4% and 42.5%, respectively, Fig. 3E), though both relative residence times are less than that of SOX2 (54%) (10). Therefore, consistent with previous studies on SOX2 and other TFs showing increased dynamics during mitosis, both HSF1-Halo and Halo-HSF2 are more dynamic in mitosis than interphase despite the difference in specific binding levels measured for the endogenous TFs via CUT&Tag (Fig. 1). These data suggest that increased TF dynamics may be inherent to the mitotic cell cycle phase and independent of site-specific binding or coating properties (see Discussion).

### Nuclear import and export activity is not sufficient to induce HSF2 coating on mitotic chromosomes

We have shown that, like SOX2, HSF2 binds site-specifically during mitosis, and that the HaloTagged TFs are similarly dynamic during mitosis. However, unlike Halo-SOX2, Halo-HSF2 is unable to coat mitotic chromosomes. We next questioned what prevents Halo-HSF2 from this coating behavior. Previous studies have shown that, despite disassembly of the nuclear envelope during mitosis, the nuclear localization signal (NLS) and the nuclear import mechanism play a role in localizing proteins to mitotic chromosomes (10, 25, 53). One possibility is that the NLS within HSF2 is insufficient to enforce coating of mitotic DNA. To test this possibility, we added three repeats of the SV40 NLS to Halo-HSF2 and to HaloTag as a control (Fig. 4A) and confirmed overexpression of each construct in mESCs (Fig. 4B). We labeled cells with JF549 and assessed protein localization during mitosis using live-cell imaging. Addition of the SV40 NLS to HaloTag induced coating on mitotic DNA (Fig. 4C), in accordance with previous reports (10). The change in coating behavior is evident in the quantification of chromatin enrichment in mitotic cells, with a mean enrichment of −0.433 for HaloTag only and a mean enrichment of 0.174 upon addition of the SV40 NLS (Fig. 4D). However, addition of the SV40 NLS to Halo-HSF2 did not cause it to localize to mitotic DNA (Fig. 4C). While the chromatin enrichment signal showed a minor increase (−1.27 vs −0.862), both constructs remain excluded from mitotic chromosomes (Fig. 4D). This data suggests that, unlike for HaloTag, the SV40 NLS is insufficient to alter Halo-HSF2 coating behavior during mitosis.

**Figure 4:**
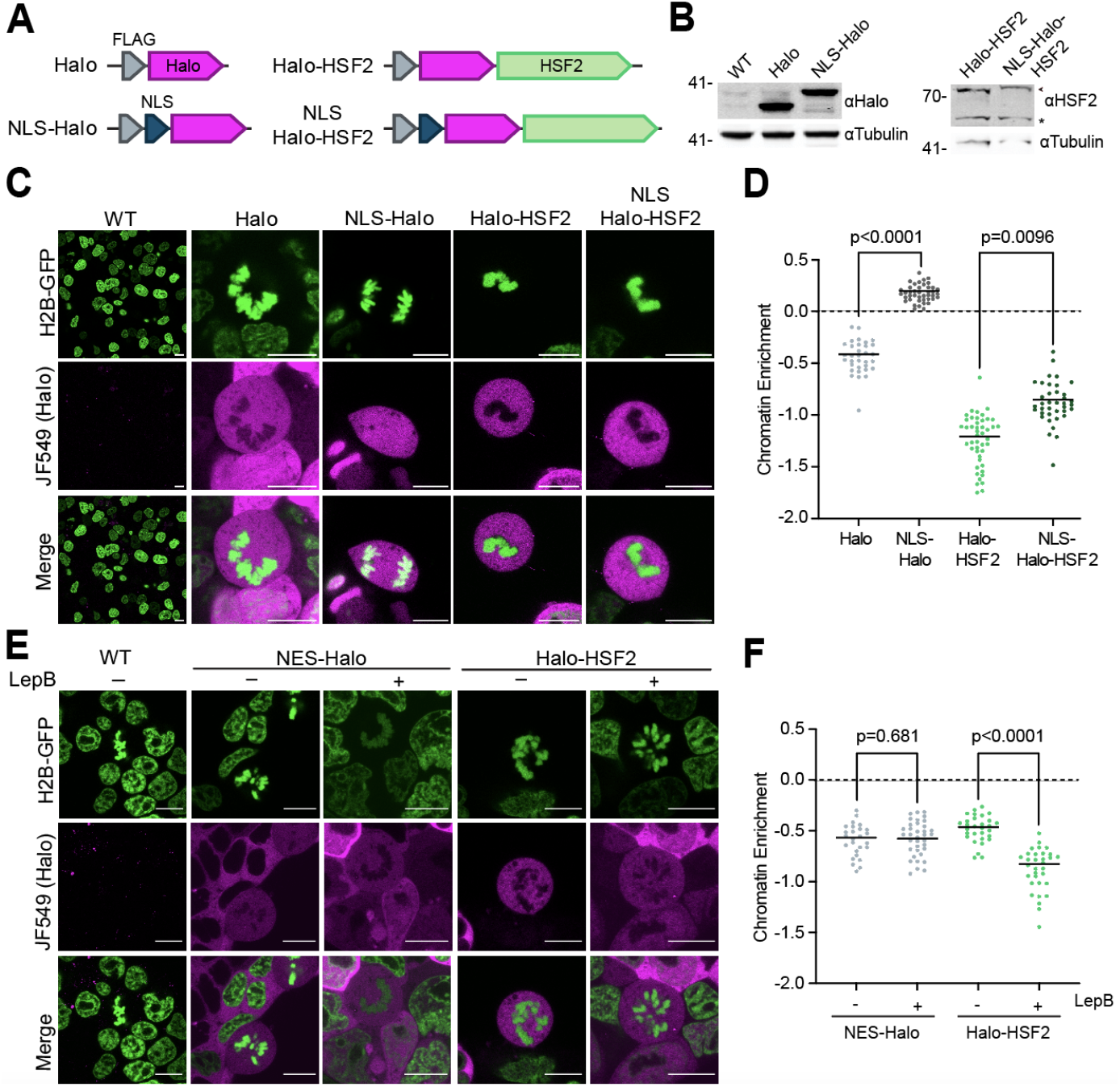
Nuclear import and export activity is not sufficient to induce HSF2 coating on mitotic chromosomes. **(A)** Schematics of stably integrated HaloTag overexpression constructs in WT JM8 cells expressing H2B-GFP, as in Fig 2A. **(B)** Protein levels of Halo and NLS-Halo constructs using α-Halo (left), and Halo-HSF2 and NLS-Halo-HSF2 constructs using α-HSF2 (right). Asterisk indicates endogenous HSF2, and arrow indicates HaloTagged protein. Tubulin is shown as a loading control. **(C, E)** Live-cell imaging of cells labeled with 200 nM JF549 dye (magenta) expressing indicated Halo constructs as well as control cells (WT). DNA is visualized by H2B-GFP overexpression (green). Scale bars are 10μm. For **(E)**, cells, were either treated with 10 ng/mL Leptomycin B (LepB) (+) or with vehicle (−) for 1h before imaging. **(D, F)** Chromatin enrichment quantification for the indicated Halo constructs (n=33 cells for Halo, 42 for NLS-Halo, 46 for Halo-HSF2, 38 for NLS-Halo-HSF2, 26 for NES-Halo control, 36 for NES-Halo +LepB, 30 for Halo-HSF2 control, and 32 for Halo-HSF2 +LepB across 3 biological replicates). Data are visualized as individual data points with mean value indicated. P-values calculated using a standard two-tailed t-test with a 95% confidence interval.

Protein localization into the nucleus is the result of two opposing mechanisms–nuclear import and export (54, 55). Since addition of the SV40 NLS is insufficient to alter Halo-HSF2 exclusion, we next examined the role of nuclear export in this process using the export inhibitor Leptomycin B (LepB) (56, 57). To validate LepB activity, we expressed HaloTag fused to the HIV-1 Rev Nuclear Export Signal (NES). This construct is cytoplasmic in untreated interphase cells, and becomes visible in the nucleus after LepB treatment, confirming drug activity (Fig. S5). We labeled mESCs expressing Halo-HSF2 with JF549 and treated the cells for 1h with LepB prior to live-cell imaging. Although the mean chromatin enrichment value for Halo-HSF2 in cells treated with LepB was much lower than those in untreated cells (−0.882 vs −0.484: Fig. 4F), overall LepB treatment had no effect on Halo-HSF2 exclusion from mitotic DNA (Fig. 4E and F). These results are consistent with the addition of the SV40 NLS described above, suggesting that neither the nuclear import or export mechanisms determine the eviction of Halo-HSF2 from chromatin during cell division.

### The heptad repeats A and B of HSF2 are necessary for exclusion from mitotic chromosomes

Our data showing that the SV40 NLS fails to affect the chromatin enrichment of Halo-HSF2 suggests that the inability of Halo-HSF2 to coat mitotic DNA may be an intrinsic property. HSF2 contains multiple functional domains (Fig. 5A): the DBD, two nuclear localization signals (NLS1 and NLS2), three heptad repeat (HR) multimerization regions (A, B, and C), a regulatory domain (RD), and a transactivation domain (TAD) (28). Apart from the DBD, the HR regions are the most relevant to DNA binding since they contribute to the trimerization of the protein, which is necessary for site-specific binding (58). HR A and B are essential for trimerization, while HR C inhibits trimerization (59).

**Figure 5:**
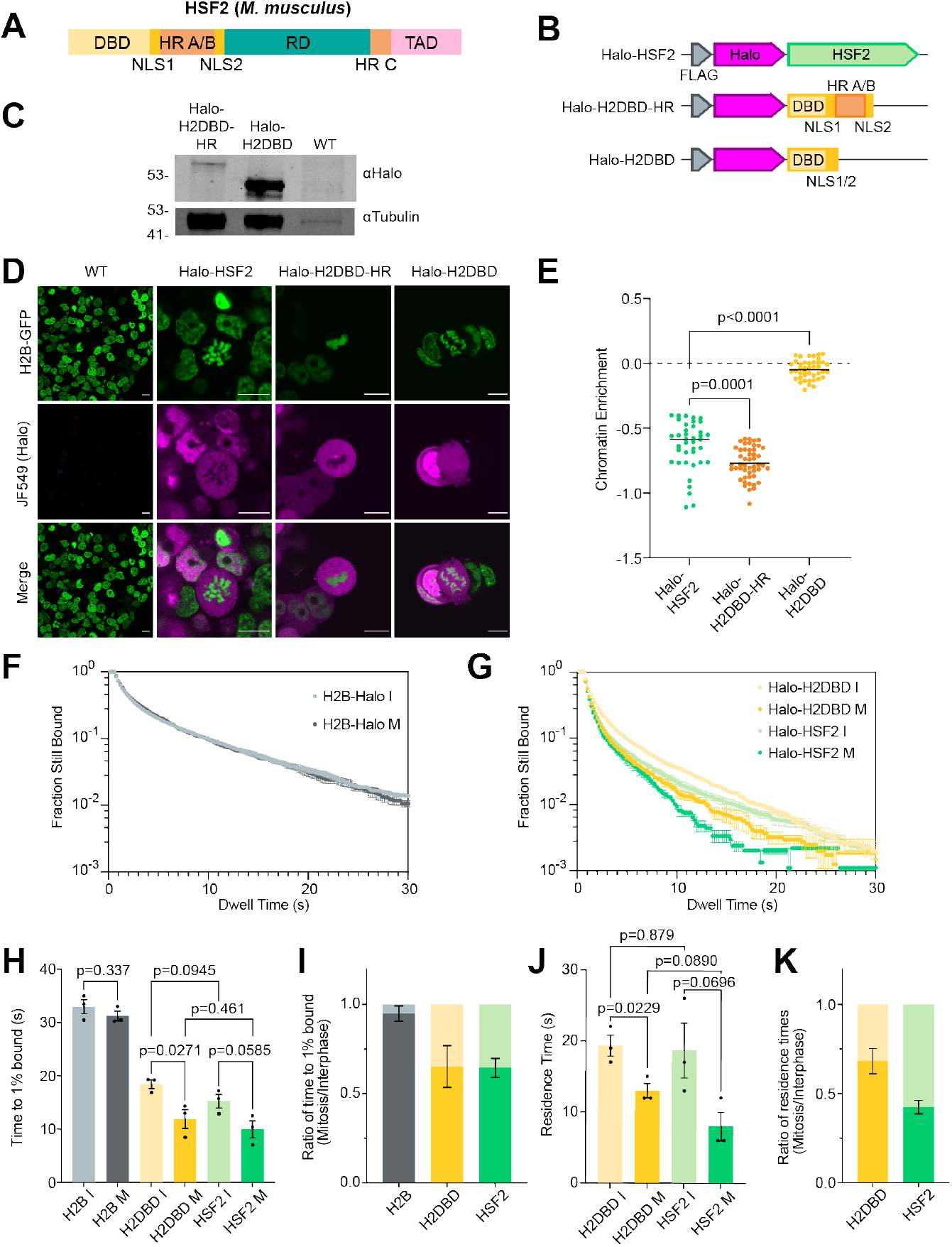
The HR region of HSF2 is necessary for exclusion from mitotic chromosomes. **(A)** Schematic showing the functional domains of HSF2. **(B)** Schematics of stably integrated HaloTag overexpression constructs in WT JM8 cells expressing H2B-GFP, as in Fig 2A. **(C)** Western blots of HaloTagged truncation constructs using α-Halo. Tubulin is shown as a loading control. **(D)** Live-cell fluorescent imaging of HaloTag-TF constructs (magenta) labeled with 200 nM JF549 dye. DNA is visualized with H2B-GFP overexpression (green). Scale bars represent 10 μm. Halo-HSF2 is presented as seen in Figure 2. **(E)** Chromatin enrichment quantification for the indicated HaloTagged TFs (n=39 cells for Halo-HSF2, 49 for Halo-H2DBD-HR, and 42 for Halo-H2DBD across 3 biological replicates). Halo-HSF2 is presented as seen in Figure 2 for comparison. **(F)** Decay curves of H2B-Halo control in interphase (I, light gray) and mitosis (M, dark gray) over time. Data depicted as mean ±SEM (n=31 cells for H2B-Halo I and 31 for H2B-Halo M across three biological replicates). **(G)** Decay curves of Halo-H2DBD (yellow), and Halo-HSF2 (green, as seen in Fig. 3A) showing the fraction of HaloTagged molecules bound to DNA in interphase (I, lighter color) and mitosis (M, darker color) over time. Data depicted as mean ±SEM (n=34 for Halo-H2DBD I, and 29 for Halo-H2DBD M across three biological replicates). **(H)** Time to 1% of total HaloTagged molecules bound to DNA for H2B-Halo control, Halo-H2DBD, and Halo-HSF2 (as in Fig. 3B) in interphase and mitosis, where each dot represents one biological replicate. Data depicted as mean ± SEM. **(I)** Ratio of the time to 1% of total HaloTagged molecules bound during mitosis relative to interphase. Data depicted as mean ±SEM. **(J)** Residence times of Halo-H2DBD and Halo-HSF2 (as in Fig. 3D) on DNA in interphase and mitosis, where each dot represents one biological replicate. Data depicted as mean ±SEM. **(K)** Ratio of mitotic residence times relative to interphase. Data depicted as mean ±SEM. All p-values are calculated using a standard two-tailed t-test with a 95% confidence interval.

To identify regions that may inhibit coating, we made two truncations of Halo-HSF2 by removing domains from the C-terminus of the protein (Fig. 5B). First, we removed the TAD, RD, and HR C of HSF2 to generate the Halo-HSF2DBD-HRA/B-NLS construct (Halo-H2DBD-HR), which contains the DBD, NLSs, and the HR A/B domains. Second, this construct was further truncated by removing HR A and B to generate the Halo-HSF2DBD-NLS construct (Halo-H2DBD). Each truncation was stably overexpressed in JM8 mESCs expressing H2B-GFP and verified with Western blots against HaloTag (Fig. 5C). The truncation constructs were imaged in live cells alongside the full-length Halo-HSF2. Removing the TAD, RD, and HR C (Halo-H2DBD-HR) did not increase chromatin enrichment (Fig. 5D,E). However, once HR A and B were removed as well, we observed a dramatic increase in chromatin coating that is visible by imaging (mean enrichment −0.770 for Halo-H2DBD-HR; −0.0480 for Halo-H2DBD) (Fig. 5D,E), though the construct is still less enriched than Halo-SOX2 (Fig. 2C). These results demonstrate that the HaloTag is not causing eviction of the protein, as the Halo-H2DBD gains the capacity to coat chromosomes. More importantly, we show that HR A and B domains are necessary for the exclusion of Halo-HSF2 from mitotic DNA, though its effects on site-specific binding remain to be seen.

We next examined if the change in coating behavior affects mitotic DNA interaction dynamics for Halo-H2DBD by performing single-molecule live-cell imaging followed by SPT. The dwell time curves for Halo-H2DBD reveal that the truncation is less dynamic in both interphase and mitosis than the fulllength protein, with Halo-HSF2 data replotted for comparison (Fig. 5G; replicate data shown in Fig. S6A). Dwell curves for the H2B-Halo control are seen in Fig. 5F. The dwell curves were fitted with a two-component exponential decay model and quantified as the time to 1% of molecules bound (Fig. 5H,I). The difference between the time to 1% bound in mitosis and interphase for Halo-H2DBD follows a similar trend as the full-length Halo-HSF2, where the interactions of the TF with DNA are more dynamic during mitosis compared to interphase (Fig. 5H). However, the difference between full-length and truncated HSF2 becomes apparent when examining the residence times and corresponding *k_off_* values of the two proteins; Halo-H2DBD consistently binds mitotic DNA for longer than Halo-HSF2 (mean=13s and 8s, respectively), despite having similar interphase residence times (mean=18.7s and 19.3s, respectively) (Fig. 5J). Reciprocally, the mean *k*_off_ in mitosis is 0.0796 s^-1^ for Halo-H2DBD and 0.138 s^-1^ for Halo-HSF2, though the mean *k*_off_ values in interphase are similar (0.052 s^-1^ for Halo-H2DBD and 0.059 s^-1^ for Halo-HSF2) (Fig. S6B). This difference is also apparent in the ratio of residence times, where Halo-HSF2 mitotic binding is 42.5% relative to interphase and Halo-H2DBD mitotic binding is 68.3% relative to interphase (Fig. 5K). This data suggests that the coating ability of Halo-H2DBD changes the mitotic residence time of the construct when compared to full-length Halo-HSF2, although the truncation remains more dynamic in mitosis than in interphase.

### The HSF2 DBD is insufficient to induce mitotic coating, in contrast to SOX TFs

We found that the HR A/B region of HSF2 counteracts the intrinsic ability of the HSF2 DBD to interact with mitotic DNA. This property contrasts sharply with SOX2 behavior where the DBD and NLS of SOX2 were shown to be the primary determinants for its mitotic chromatin association (10). Three point mutations in the SOX2 DBD (M47G, F50G, and M51G) were enough to abolish specific DNA-binding ability (50), and resulted in the exclusion of SOX2 from mitotic DNA (10). Additionally, the SOX2 DBD is able to coat mitotic chromatin in isolation (10). Thus, to complement our studies with the HSFs, we investigated the importance of the DBD for mitotic chromatin coating within the SOX TF family. SOX2 and SOX13 share 55% DBD sequence similarity (Fig. S6C), but SOX2 is enriched on mitotic DNA (10, 18) whereas SOX13 is excluded (18). Given the importance of the DBD for the mitotic coating of SOX2, we hypothesized that exchanging the DBDs between SOX2 and SOX13 could reverse their mitotic enrichment. We generated and expressed four HaloTagged TF constructs: full-length Halo-SOX2 (as seen in previous figures) and Halo-SOX13, and two constructs with the DBDs exchanged (Fig. 6A,B). As described previously (10, 18), Halo-SOX2 is strongly enriched on mitotic chromatin while Halo-SOX13 is clearly excluded (Fig. 6C,D). When the SOX2 DBD is replaced with that of SOX13 (Halo-S2S13DBD), the construct is evicted from mitotic DNA (mean enrichment −0.383) (Fig. 6C,D). Accordingly, when the SOX13 DBD is replaced with that of SOX2 (Halo-S13S2DBD), the construct coats to a similar degree as full-length Halo-SOX2 (mean enrichment 0.861 and 0.790, respectively) (Fig. 6C,D). Thus, the SOX2 DBD is both necessary and sufficient to induce mitotic chromatin coating.

**Figure 6:**
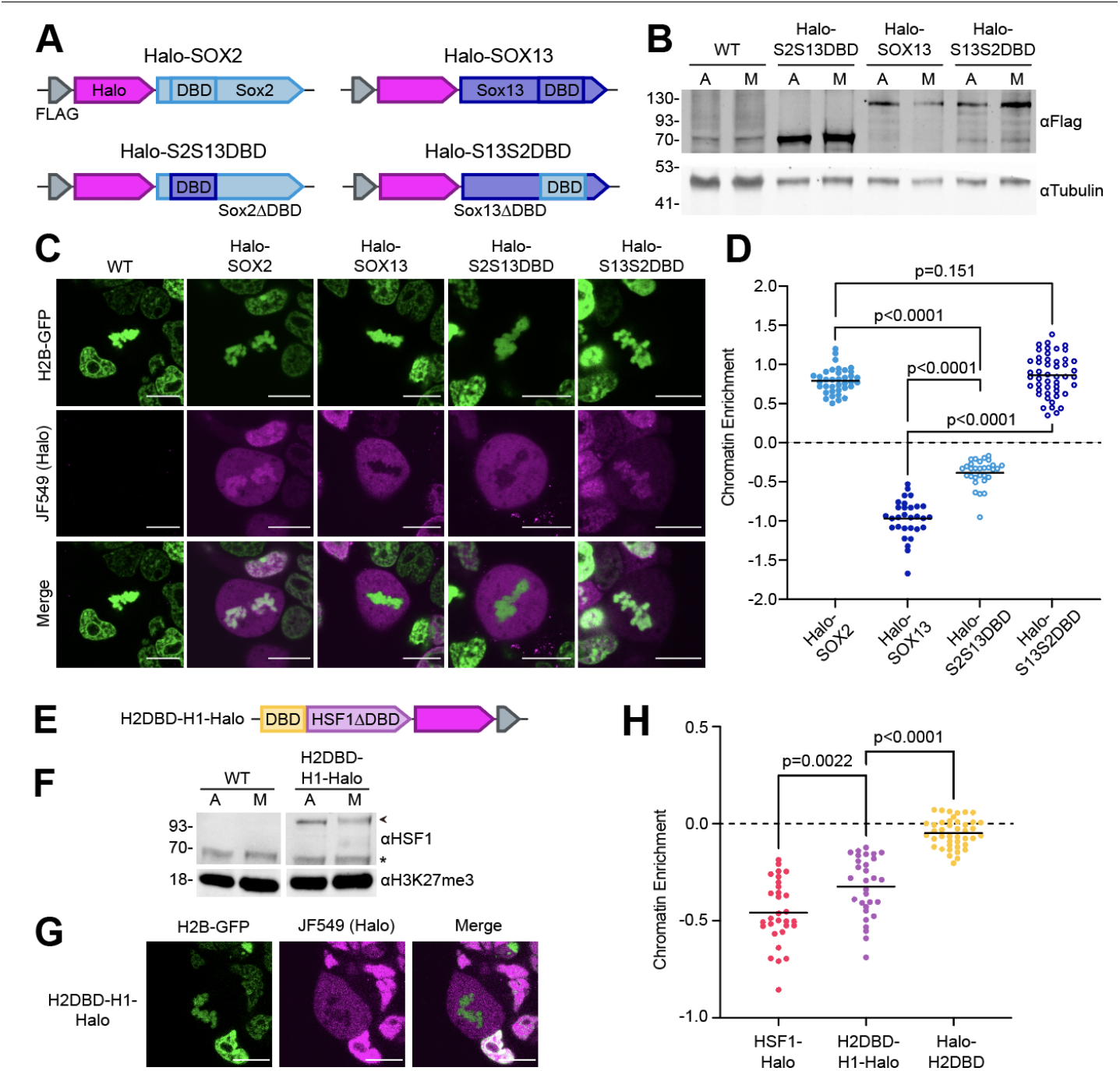
The HSF2 DBD is not sufficient to induce mitotic coating, in contrast to SOX TFs. **(A)** Schematics of stably integrated HaloTag overexpression constructs in WT JM8 cells expressing H2B-GFP, as in Fig 2A. **(B)** Western blots of each HaloTag-TF construct in asynchronous (A) and mitotic (M) cells using α-Flag. Tubulin is shown as a loading control. Mitotic cell populations were synchronized with 50 ng/mL nocodazole for 6 hours before collecting cells via shake-off. **(C)** Live-cell fluorescent imaging of HaloTag-TF constructs (magenta) labeled with 200 nM JF549 dye. DNA is visualized with H2B-GFP overexpression (green). Scale bars represent 10 μm. **(D)** Chromatin enrichment quantification for the indicated HaloTagged TFs (n=37 cells for Halo-Sox2, 30 for Halo-Sox13, 30 for Halo-S2S13DBD, and 48 for S13S2DBD across 3 biological replicates). P-values calculated using a standard two-tailed t-test with a 95% confidence interval. **(E)** Schematic of the HSF2DBD-HSF1-Halo construct (H2DBD-H1-Halo) which was stably integrated into WT JM8 cells expressing H2B-GFP. **(F)** Western blot of H2DBD-H1-Halo in asynchronous (A) and mitotic (M) cells using α-HSF1. H3K27me3 shown as a loading control. Mitotic cell populations were synchronized as in (B). **(G)** Live-cell fluorescent imaging as in (C). **(H)** Chromatin enrichment quantification for H2BD-H1-Halo compared to full-length HSF1-Halo (data as shown in Fig. 2E) and Halo-H2DBD (data as shown in Fig. 5E). n=30 cells for Halo-H2DBD-H1 across 3 biological replicates. P-values calculated using a standard two-tailed t-test with a 95% confidence interval.

Knowing that the HSF2 DBD alone is able to coat mitotic chromosomes (Fig. 5), we next asked whether the HSF2 DBD is sufficient to induce the mitotic coating of HSF1-Halo. To address this question, we generated and expressed the HSF2DBD-HSF1-Halo construct (H2DBD-H1-Halo), in which the DBD of HSF1 is replaced with the HSF2 DBD (Fig. 6E,F). Live-cell imaging of H2DBD-H1-Halo revealed clear exclusion from mitotic DNA as also seen for HSF1-Halo (Fig. 6G,H). Though replacement of the HSF1 DBD with HSF2 DBD did increase enrichment (mean enrichment −0.458 and −0.325, respectively), this effect is moderate in comparison to the effects of removing HR A/B from HSF2 DBD (Fig. 5D,E). Additionally, H2DBD-H1-Halo is still clearly excluded, while Halo-H2DBD alone was visibly enriched (Fig. 5D). Therefore, the HFS2 DBD is not sufficient to induce mitotic coating of HSF1-Halo. This data further suggests that parts of the non-DBD domains in HSF1 promote exclusion from mitotic DNA in a similar manner as the HSF2 HR. Thus, while the mitotic behavior of TFs like SOX2 and SOX13 is driven by the DBD, exclusion of HSF1-Halo and Halo-HSF2 from mitotic DNA is directed by non-DBD domains.

## Discussion

In this paper, we investigate the mitotic DNA interactions of two related TFs: HSF1 and HSF2. Though the DBDs of these TFs are highly conserved, HSF2 maintains site-specific binding to mitotic DNA whereas HSF1 does not. Despite this difference in specific binding, imaging analyses show surprisingly similar behaviors when these factors interact with mitotic DNA. Neither HaloTagged HSF1 nor HSF2 coat mitotic chromosomes as seen with live-cell imaging, and both HSF1-Halo and Halo-HSF2 interact with mitotic DNA more dynamically than with interphase chromatin. Addition of the SV40 NLS to Halo-HSF2 or inhibiting active nuclear export failed to alter the interactions of Halo-HSF2 with mitotic DNA, suggesting that these pathways do not influence coating behavior for HSF2. Instead, the isolated HSF2 DBD is able to coat mitotic chromosomes, and this coating is directly inhibited by the inclusion of HR A and B domains in this construct. Furthermore, we observed that Halo-H2DBD has a longer residence time on mitotic DNA than Halo-HSF2, suggesting that at least part of the dynamic mitotic DNA-HSF2 interactions are due to the non-DBD regions. Though able to coat on its own, the HSF2 DBD is unable to induce mitotic coating of HSF1, indicating that the non-DBD domains of the HSFs not only contribute to increased dynamics but also dictate their exclusion from mitotic DNA.

We have shown that HSF2 can bind site-specifically during mitosis but remains generally excluded from mitotic chromosomes as visualized by imaging, demonstrating that site-specific binding and global mitotic coating are distinct abilities that can be measured with different methods. These results bring into question the true nature of DNA-TF interactions in mitosis. Currently, studies using live-cell imaging approaches have shown that many TFs coat with mitotic DNA (10, 14–16, 18), but precisely how coating translates to specific binding has been in question (11), largely because the methods used to measure specific binding, such as ChIP-seq, are reliant on fixation that alters DNA-TF interactions in mitosis (10). In some cases, coating of TFs on mitotic DNA corresponded to site-specific binding. For instance, the pluripotency transcription factors ESRRB (11, 16) and SOX2 (Fig. 1 and 2), both have a strong capacity to coat mitotic DNA, and profiling their mitotic DNA interactions using genomic approaches reveals hundreds of occupied loci. However, some evidence also suggests that the coating of mitotic DNA by TFs does not directly relate to specific DNA-binding events. For example, a mutant version of FOXA1 that cannot recognize and bind to specific target sites was largely retained on mitotic chromosomes as visualized by live imaging, despite the loss of binding on specific loci (15). Therefore, previous studies have shown three modes of TF-mitotic DNA interactions: coating with site-specific binding, coating without site-specific binding, and neither coating nor binding site-specifically. Our study reveals a fourth mode: HSF2 cannot coat mitotic chromosomes but can bind specifically to target loci (Fig. 1 and 2). It remains to be seen how many TFs fit into each of these categories of TF-mitotic DNA interactions.

Regardless of which mitotic DNA-TF interaction category they fit in, single-molecule live-cell imaging and SPT revealed similar DNA-TF interaction dynamics between HSF1-Halo, Halo-HSF2, and Halo-H2BDD in mitosis and interphase (Fig. 3A-C and Fig. 5H,I). All of these constructs interacted with DNA more dynamically in mitosis than in interphase. This consistent pattern of increased DNA interaction dynamics during mitosis has also been shown for various other TFs using SPT or fluorescence recovery after photobleaching, including SOX2 (10), FOXA1 (15, 18), CDX2 (18), HMGB2 (18), POU5F1 (18), TEAD1 (18), and others. The mechanisms underlying these increased dynamics for TFs examined thus far are not fully understood. It is possible that the drastic changes to the cellular landscape during mitosis contributes to these increased DNA-TF dynamics. For instance, chromosomes are globally reorganized and condensed during mitosis, which may affect the association rates and target search mechanism used by TFs. In interphase, TFs use a mechanism known as facilitated diffusion to slide across DNA scanning nucleotides (60, 61). The DBD of TFs has a non-specific affinity for DNA and a high specific affinity for its binding motif (62), though only a fraction of potential TF binding motifs serve as highly occupied targets (63). The tight packing of nucleosomes in mitotic chromatin may disrupt this target search, leading to fewer interactions with potential binding sites and thus increased *k*_off_. Another global change in mitosis is the dissolution of the nuclear membrane, which increases the space in which a TF can diffuse and may also impact association rates and target search. Furthermore, binding of many TFs is stabilized by cooperativity with other proteins (64–67), and loss of these protein-protein interactions during mitosis, due to the global and massive decrease in transcriptional activity, may also contribute to the increased *k_off_* observed for TFs at this time (Fig. S4C, S6B). Surprisingly, our SPT data for the full-length versus DBD truncation of HSF2 suggests that at least part of the DNA-TF dynamics may be inherent to specific TF domains. The relative residence times for Halo-HSF2 in mitosis is 42.5% relative to interphase, and is increased to 68.3% for Halo-H2DBD (Fig. 3E, Fig. 5K), indicating that on average the DBD-only molecules interact with mitotic DNA longer than full-length HSF2. Interestingly, this number is even higher than for SOX2, the residence time of which is 54% in mitosis relative to interphase (10). The domains removed from the HSF2 DBD truncation include the trimerization domains (HR A, B, and C), the regulatory domain, and the transactivation domain; how these domains specifically contribute to increased mitotic DNA-TF interaction dynamics remains to be seen.

What drives the differences in behavior of highly similar TFs during mitosis? Previous studies suggested that the DBD is one of the major determinants of mitotic DNA coating (10, 25, 68), Indeed, SOX2 DBD alone is able to coat mitotic DNA (10), and our results show that the SOX2 DBD can induce mitotic coating of SOX13 (Fig. 6 C,D). However, at least in the case of the HSFs, we show that the DBD is not the dominant driver of mitotic DNA interactions. In fact, even though the HSF2 DBD alone can coat mitotic chromosomes, it is unable to induce mitotic DNA coating not only of full-length HSF2, but also of the non-DBD domains of HSF1. Therefore, the non-DBD regions of HSF2, perhaps the trimerization domain, not only contribute to DNA-TF interaction dynamics, but ultimately drive the mode of HSF2 behavior during mitosis (i.e. not coating but binding site-specifically). Both HSF1 and HSF2 generally bind to DNA as trimers (69, 70), which form a stable DNA-enveloping structure (71, 72). In vitro studies have revealed that monomeric HSF1 is able to bind DNA, albeit at lower affinity than trimerized protein (73). The equilibrium dissociation constant (K_D_) of monomeric HSF1 was ~10 times higher than that of the trimer, whereas trimerization deficient HSF1 with the HR domains excised had ~70 times higher K_D_. Our SPT results show that the Halo-H2DBD, which also lacks the trimerization HR domains, has only slightly lower *k_off_* rates (longer residence times) on mitotic chromosomes compared to the full length HSF2 (Fig. S6B). Assuming that the binding affinity of HSF2 trimers is similar to that of HSF1, and that K_D_ = *k_off_/k_on_* (59), *k_on_* for the truncated H2DBD protein would be at least 20x decreased compared to the full length HSF2. The decreased *k_on_* means a much slower rate of finding specific target sites, which could signify increased time spent in diffusion and/or target search and non-specific DNA associations (75). The change in coating behavior that we observed for Halo-H2DBD (Fig. 5D-E) would be consistent with increased non-specific DNA associations. Taken together, our findings highlight the importance of multimerization domains like the HR in mitotic DNA association of the HSFs. Since many TFs act as dimers or trimers, such multimerization capacity could determine the mode of TF-mitotic DNA interactions for these TFs.

## Materials and Methods

### Cell Culture

Mouse ES cells (JM8.N4, RRID: CVCL_J962) were used for all experiments and obtained as previously described (14). ES cells were cultured on 0.1% gelatin-coated plates in ESC DMEM (Corning 10101CV, contains 4.5 g/L glucose, 110 mg/L sodium pyruvate, glutagro, and 15 mg/L phenol red) with 15% FBS (HyClone), 0.1 mM MEM non-essential amino acids (Gibco), 2 mM L-Glutamine (Gibco), 0.1 mM 2-mercaptoethanol (Sigma), 1X Penicillin Streptomycin solution (Corning), and 1000 units/ml of ESGRO LIF (Chem-icon). ESCs were fed daily, cultured at 37 °C in a 5% CO_2_ incubator, and passaged every two days by trypsinization. For Leptomycin B treatment, cells were treated with 10 ng/mL Leptomycin B (Sigma) at 37 °C in a 5% CO_2_ incubator for 1 hour. For all imaging experiments, cells were imaged in DMEM without phenol red (Gibco 31053028, contains 4.5 g/L glucose) with 110 mg/L sodium pyruvate (Gibco) and all other components previously listed.

### Mitotic Synchronization

mESCs were treated with 50 ng/mL nocodazole (Sigma) at 37 °C in a 5% CO_2_ incubator for 6 hours. After nocodazole removal, cells were washed with 1X PBS and mitotic cells were collected via shake-off. Cells were fixed in 70% ethanol and purity of the mitotic fraction (mitotic index) was monitored by flow cytometry and cell counting. For flow cytometry, asynchronous and shake-off cells were stained with propidium iodide (20 μg/ml; Sigma-Aldrich) and analyzed on the FACS Canto system (BD Bioscience). The flow cytometry profiles were analyzed using FlowJo software. Live cells were gated and the histogram function was plotted. For cell counting, cells were permeabilized with 0.1% Triton-X, stained with 300 nM DAPI (Invitrogen), and images were acquired using a microscope (Leica DMI6000 B). Mitotic cells were identified based on the presence and morphology of condensed mitotic chromosomes and quantified as percent of total cells within the image field.

### Cloning

To generate HaloTagged TFs, the coding sequence of each TF was cloned into a Piggybac vector containing either a 3X-Flag-Halo-TEV construct upstream of a multiple cloning site (used for all N-terminal HaloTagged TF constructs), or a TEV-Halo-1X-Flag construct downstream of a multiple cloning site (used for all C-terminal TF-Halo constructs). All primers used for cloning are listed in Supplementary Table 1. All HaloTagged TF plasmids are deposited on Addgene.

### Transfection

Mouse ES cells were grown to ~50% confluency on 0.1% gelatin-coated 6-well plates at 37 °C in a 5% CO_2_ incubator. Stably expressing cell-lines were generated by transfecting 1 μg of the HaloTagged TF Piggybac construct together with 1 μg of the Super Piggybac transposase plasmid (gifted from the Tjian Lab) using Lipofectamine 2000 (Invitrogen) according to the manufacturer’s instructions. After 24 hours, 500 μg/mL G418 (Fisher) was added. Cell media containing antibiotics was refreshed daily until all negative control cells were dead. A similar method was used to stably express H2B-GFP under 0.6 μg/mL Puromycin (ThermoFisher).

### Generation of HSF1 and HSF2 knock-out cells with CRISPR-Cas9

Two sets of guide RNAs (gRNAs) targeting the entire locus of HSF1 or HSF2 respectively were designed using Benchling CRISPR Guide RNA Design tool (https://www.benchling.com/crispr) and cloned into the px458 gRNA expression plasmid. Mouse ES cells were transfected with 1 μg of plasmid containing Cas9 and gRNAs using Lipofectamine 2000 (Invitrogen) according to the manufacturer’s instructions. Cells were sorted two days after transfection, and 20000 transfected-positive cells were plated onto a 15 cm tissue culture plate. One week after plating, individual colonies were picked up and trypsinized. 80% of each cell colony was used for genotyping, while 20% of each cell colony was grown for maintenance. For genotyping, cells were lysed using DirectPCR lysis reagent (Viagen Biotech) according to the manufacturer protocol and lysates were used in a screening PCR to identify edited cells. Identified clones were further validated using Sanger sequencing and Western blotting. A list of gRNAs and primers used for gene editing are in Supplementary Table 2.

### Live-cell Imaging

Mouse ES cells were grown on 0.1% gelatin-coated chambered coverslips (Ibidi, μ-Slide 4 Well 80426 or 2 Well 80286) at 37 °C in a 5% CO_2_ incubator for 24 hours. Cells were labeled with 200 nM JF549 (Lavis Lab) for 30 minutes at 37 °C in a 5% CO_2_ incubator and washed 3 times with 1X PBS for 5 minutes each. Cells were imaged with a Leica SP8 confocal microscope at 37 °C in ESC DMEM without phenol red. Chromatin enrichment quantification was performed with ImageJ as follows: the mean HaloTag fluorescence intensity was calculated in the area of the mitotic chromatin and across the whole mitotic cell. The chromatin enrichment score was calculated by taking log_2_ of the chromatin mean divided by the whole cell mean.

### Western Blot

Mouse ES cells were cultured to ~90% confluency on 0.1% gelatin-coated plates at 37 °C in a 5% CO_2_ incubator. After indicated treatments, cells were washed with 1X PBS, trypsinized, and pelleted by centrifuging at 600 g. Cell pellets were lysed in lysis buffer (150 mM NaCl, 50 mM HEPES pH 7.6, 1 mM EDTA, 2 mM MgCl2, 1% Triton X, 10% Glycerol, 1X Roche cOmplete EDTA-free Protease Inhibitor, 5 mM PMSF) for 15 minutes on ice. Lysates were pelleted at 4 °C for 15 minutes at 16,000 g. The supernatants were saved, mixed with 4X Laemmli buffer, and boiled at 95 °C for 5 minutes. Proteins were separated via 7, 8, 9, or 10% SDS-PAGE and wet transferred to Nitrocellulose membranes (VWR, CA10061-084) at 4 °C. Membranes were blocked with blocking buffer [1X PBS, 0.1% Tween (Fisher)] containing either 5% non-fat milk powder or 3% BSA (Sigma-Aldrich) at room temperature for 40 minutes. Membranes were incubated in α-HSF1 1:1000 (Abcam, ab2923), α-HSF2 1:1000 (32), α-H3K27me3 1:7000 (Cell Signaling Technologies, C36B11), α-Tubulin 1:7000 (Abcam, ab6046), α-SOX2 1:1000 (Cedarline, 39844), α-HaloTag 1:1000 (Promega, G9211), or α-Flag 1:5000 (Sigma-Aldrich, F3165) overnight at 4°C. Membranes were washed in 1X PBS with 0.1% Tween 3 times for 5 minutes each at room temperature. Membranes were then incubated in IRDye 800CW Goat anti-mouse (Cedarlane, 926-32210) or IRDye 800CW Goat anti-rabbit (Cedarlane, 925-32211) secondary antibodies (1:20000) at room temperature for 45 minutes. Finally, membranes were washed in 1X PBS with 0.1% Tween 3 times for 5 minutes each before scanning using the ChemiDoc™ MP Imaging System (Bio-Rad). Note: membranes blotted with α-HSF2 were boiled for 10 minutes prior to blocking (33).

### Single Particle Tracking: Slow Tracking

Cells were grown on gelatin-coated 35mm glass-bottomed dishes (Ibidi, μ-Dish 35mm, high Glass Bottom 81158) at 37 °C in a 5% CO_2_ incubator for 24 hours. Cells were labeled with 25 pM (H2B-Halo control) or 40 pM (HaloTagged TFs) JF549 (Lavis Lab) for 30 minutes at 37 °C in a 5% CO_2_ incubator and washed 3 times with ESC media without phenol-red for 5 minutes each. Cells were imaged in ESC media without phenol-red. Imaging was conducted on a custom-built 3i (Intelligent Imaging Innovations) microscope equipped with a Alpha Plan-Apochromat 100x/1.46 NA oil-immersion TIRF M27 objective, EM-CCD camera (Andor iXon Ultra 897), a Zeiss Definite Focus 2 system and a motorized mirror to achieve HiLo-illumination. The customized laser launch includes 405 nm (350 mW), 488 nm (300 mW), 561 nm (1 W) and 640 nm (1 W) lasers. A multi-band dichroic (405 nm/488 nm/561 nm/633 nm quadband bandpass filter) was used to reflect a 561 nm laser into the objective and emission light was filtered using a bandpass emission filter. The laser intensity was controlled using an acousto-optic transmission filter. A low constant laser intensity was used to minimize photobleaching. Images were collected at a frame rate of 5 Hz for a total of 1000 frames. Each Halo-tagged line was imaged in 3 biological replicates of 6-12 cells.

HaloTagged TF particles were identified using SLIMfast (34), a custom-written MATLAB implementation of the MTT algorithm (35), using the following algorithm settings: localization error: 10^−6.25^; exposure time: 200 ms; deflation loops: 3; number of gaps allowed: 1; maximum number of competitors: 5; maximal expected diffusion constant (μm^2^/s): 0.5. The residence times of HaloTagged TFs were determined using custom scripts as previously described (10, 14). Briefly, we quantified the dwell time of each HaloTagged TF molecule and generated a survival curve for bound HaloTagged TFs as a function of time. A two-exponential function was fitted to the survival curve to determine apparent *k*_off_ rates of HaloTagged TFs and was used to calculate the time to 1% bound shown in Figures 3B, 3C, 5H, and 5I. Photobleaching correction was performed by subtracting the apparent *k*_off_ of H2B-Halo from the apparent *k*_off_ of the HaloTagged TFs, and residence time was determined by taking the inverse of the photobleach-corrected *k*_off_ for each HaloTagged TF.

### CUT&Tag

CUT&Tag assay was performed as previously described (31). Briefly, 100,000 mESCs were used per sample and were bound to Concanavalin A-coated beads (Cedarlane Labs BP531-3ML). Cryopreserved *Drosophila melanogaster* S2 cells were spiked in at a 10% concentration (10,000 S2 per 100,000 mESCs). Cells attached to beads were incubated at 4 °C overnight with primary antibodies. Epicypher pAG-Tn5 (EpiCypher EP151117) was used at 1:20 final concentration and tagmentation was performed for 1 hour in a 37 °C water bath. DNA fragments were solubilized in STOP solution by overnight incubation in a 37 °C water bath. Library preparation was also performed as in the original protocol.

The following primary antibodies were used for CUT&Tag at 1:100 dilution: α-HSF1 (Abcam, ab2923), α-HSF2 (32), α-H3K27me3 (Cell Signaling Technologies C36B11), α-SOX2 (Cedarlane 39844), Rabbit α-Mouse IgG (Abcam ab46540). For the secondary antibody, Guinea Pig α-Rabbit IgG (Antibodies-Online ABIN101961) at 1:100 dilution was used.

### CUT&Tag Analysis

Reads were mapped on mm10 genome build using Bowtie2 version 2.4.2 (36) with the following parameters: --local --very-sensitive-local --no-unal --no-mixed --no-discordant --phred33 -I 10 -X 2000. PCR duplicate reads were kept as these sites may represent real sites from adapter insertion from Tn5 as described previously (37). To quantify reads from S2 spiked in cells, reads were mapped on dm6 genome build using Bowtie2 as above with additional parameters: --no-overlap --no-dovetail. The number of reads aligned to the *D. melanogaster* genome per sample was used to calculate scaling values, where an arbitrary constant (100,000) was divided by the number of fly reads and normalized such that the asynchronous sample was set to 1. After alignment, BAM files were converted to bigwig files scaled with the calculated scaling values (Supplementary Table 3) using deepTools and visualized as gene tracks using IGV (38). Heatmap analyses were performed using DeepTools and BedTools suite (39, 40). ComputeMatrix from deepTools was done using binsize 10. Peak calling for HSF1 and HSF2 CUT&Tag was performed using the SECAR suite (41) with options: “non” and “stringent”, using IgG signal as the control. Bedgraph files for SECAR were prepared with the code provided with the SECAR suite. The peaks identified in asynchronous and mitotic samples were combined and filtered for unique sites to generate a BED file of binding sites. Read counts for scatter plots were obtained for each TF from BAM files using the bedtools multicov command against a BED file with binding sites for each of TFs. Read counts were then normalized to the *D. melanogaster* scaling values and plotted using the ggplot2 suite in R. Pearson correlation analysis was performed using deepTools multBamSummary with the BED option. The Pearson correlation coefficients were plotted as a heatmap using the plotCorrelation command with the remove outliers and skip zeros options.

### Putative Binding Sites Analysis

Primary binding motifs for mouse HSF1, HSF2, and SOX2 were obtained from HOCOMOCO database (42) as Position Weight Matrices (PWMs). PWMs were then used as an input motif in the FIMO tool (43) to identify putative binding sites in the mouse (mm10) genome with p-value threshold set below 1×10^-4^.

### Gene Ontology Analysis (GEO)

Peaks identified using the SECAR suite were annotated using the HOMER annotatePeaks function (44). Entrez ID of the nearest genes were used for GEO analysis using the DAVID suite (45, 46).

## Data and Code Availability

The CUT&Tag datasets generated in this study have been deposited to Gene Expression Omnibus (https://www.ncbi.nlm.nih.gov/geo/), and are available as raw and processed files through accession numbers GSE214219. Plasmid constructs are deposited on Addgene.

## Acknowledgements

We thank Dr. Steven Henikoff for generously providing the pA-Tn5 enzyme. We thank R. Vander Werff and T. Stach (BRC-seq, UBC) for Illumina sequencing. This work was supported by Life Sciences Institute Cores (LSI Imaging and ubcFLOW), supported by the UBC GREx Biological Resilience Initiative. For insightful comments on the manuscript, we thank Thomas Nguyen and Drs. Annie Ciernia, Ethan Greenblatt, and Calvin Yip. S.S.T is a Wall Scholar at the Peter Wall Institute, and a Michael Smith Foundation for Health Research Scholar.

## Author contributions

Conceptualization: RMP, MAB, SST

Methodology: RMP, MAB, JS, JEM, JZJK

Investigation: RMP, MAB, JS, JEM, JZJK

Visualization: RMP, MAB, JS, JEM

Funding acquisition: RMP, MAB, JS, SST

Supervision: SST

Writing – original draft: RMP, MAB, SST

Writing – review and editing: RMP, MAB, JS, JEM, JZJK, SST

## Supplementary Figures

**Supplemental Figure 1:**
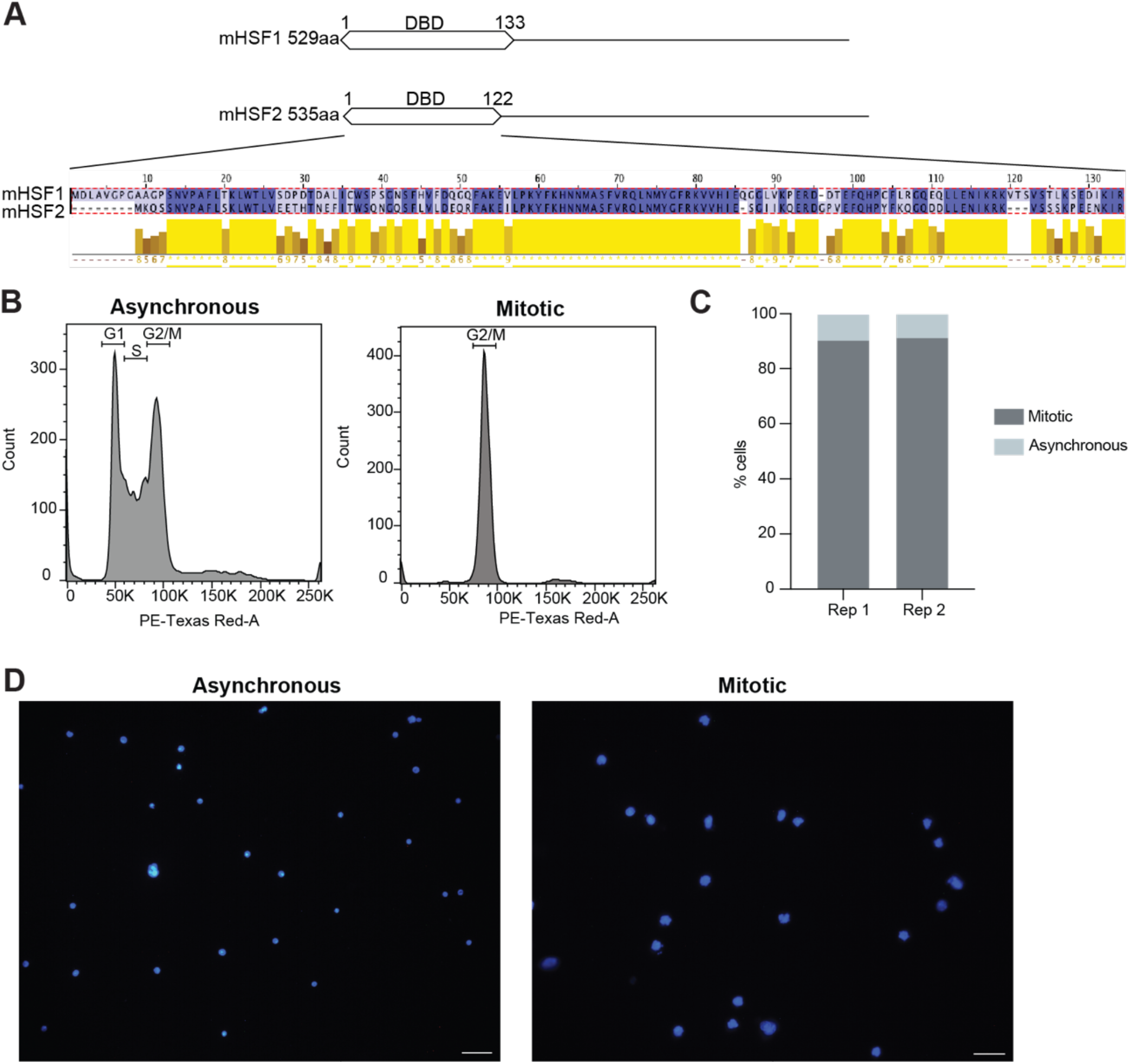
Efficiency of mitotic enrichment. **(A)** HSF1 DBD and HSF2 DBD sequence alignment (blue) and scoring matrices (yellow). Alignment produced with Jalview. **(B)** Representative histograms of asynchronous and mitotic cells based on the DNA content. **(C)** Mitotic indexes of biological replicates used as mitotic fractions for CUT&Tag assay. **(D)** Representative images of DAPI-stained cells used for calculation of mitotic index in panel B. Scale bar represents 100 μm.

**Supplemental Figure 2:**
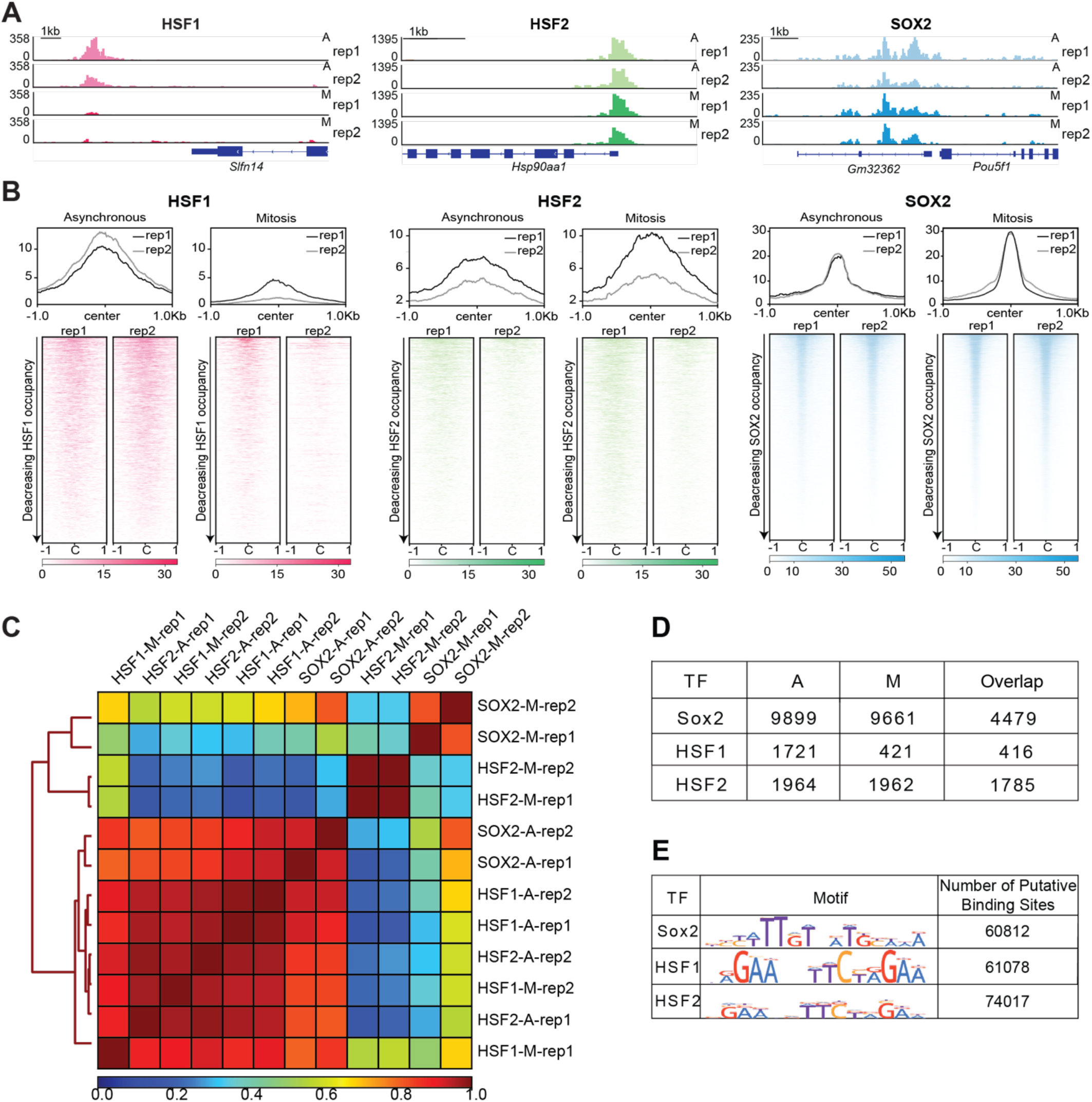
Replicate analysis of CUT&Tag data and analysis of TF binding in asynchronous and mitotic cells. **(A)** Gene browser tracks of biological replicates for HSF1 over *Slnf14* locus, HSF2 over *Hsp90aa1* locus and SOX2 over *Pou5f1* locus in asynchronous (A) and mitotic (M) cells. **(B)** Genome-wide average plots (top) and heatmaps (bottom) of biological replicates of HSF1 (left), HSF2 (middle), and SOX2 (right) CUT&Tag in asynchronous (A) and mitotic (M) cells. CUT&Tag signal was calculated in a 2 kb window surrounding binding sites of the respective TF. For heatmaps, binding sites were ordered by decreasing occupancy of the given TF. **(C)** Pearson correlation analysis of HSF1, HSF2, and SOX2 biological replicates in asynchronous (A) and mitotic (M) cells. Average score is based on coverage for regions that cover HSF1, HSF2 and SOX2 binding loci. **(D)** Number of SOX2, HSF1, and HSF2 target loci bound in asynchronous (A) and mitotic (M) cells and number of overlapping loci between A and M. **(E)** Number of putative binding sites for SOX2, HSF1, and HSF2 throughout the mouse genome. Binding motifs used for search of potential binding sites are shown for each TF.

**Supplemental Figure 3:**
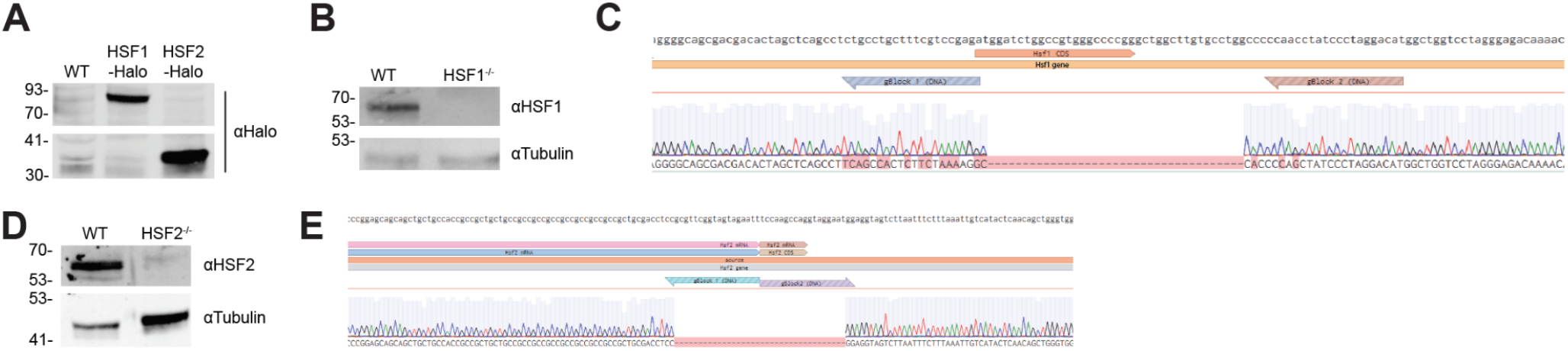
C-terminal HaloTagging of HSF2, and HSF knock-out verification. **(A)** Western blot of overexpressed HSF1-Halo and HSF2-Halo in JM8 cells using α-Halo. C-terminal tagging of HSF2 resulted in truncation of the protein. **(B)** Western blot of WT and HSF1^-/-^ JM8 cells using α-HSF1. Tubulin is shown as a loading control. **(C)** Sanger sequencing trace showing region of *HSF1* gene deleted in HSF1^-/-^ JM8 cells. **(D)** Western blot of WT and HSF2^-/-^ JM8 cells using α-HSF2. Tubulin is shown as a loading control. **(E)** Sanger sequencing trace showing region of *HSF2* gene deleted in HSF2^-/-^ JM8 cells.

**Supplemental Figure 4:**
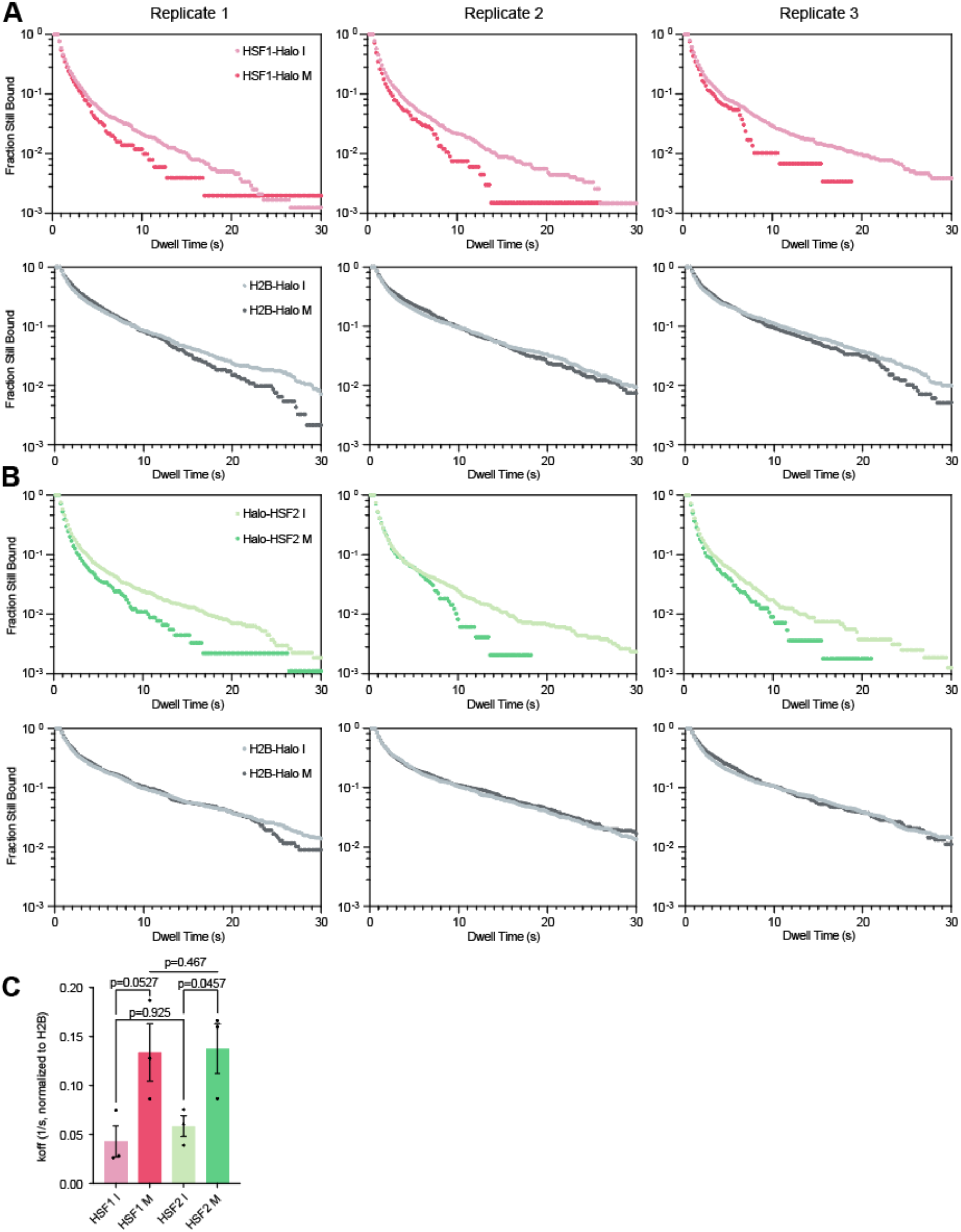
Replicate dwell curves for HSF1-Halo and Halo-HSF2 SPT. **(A)** Decay curves for each biological replicate of HSF1-Halo (red, top) and the associated H2B-Halo replicate (gray, bottom) representing the fraction of HaloTagged molecules bound to DNA in interphase (I, lighter color) and mitosis (M, darker color) over time. Technical replicates: n=19 cells for H2B-Halo I Rep1, 11 for H2B-Halo I Rep2, 10 for H2B-Halo I Rep3, 9 for H2B-Halo M Rep1, 11 for H2B-Halo M Rep2, 10 for H2B-Halo M Rep3, 12 for HSF1-Halo I Rep1, 12 for HSF1-Halo I Rep2, 10 for HSF1-Halo I Rep3, 5 for HSF1-Halo M Rep1, 10 for HSF1-Halo M Rep2, and 8 for HSF1-Halo M Rep3. **(B)** Decay curves for each biological replicate of Halo-HSF2 (green, top) and the associated H2B-Halo replicate (gray, bottom) representing the fraction of HaloTagged molecules bound to DNA in interphase (I, lighter color) and mitosis (M, darker color) over time. Technical replicates: n=12 cells for H2B-Halo I Rep1, 9 for H2B-Halo I Rep2, 10 for H2B-Halo I Rep3, 9 for H2B-Halo M Rep1, 11 for H2B -Halo M Rep2, 10 for H2B -Halo M Rep3, 11 for Halo-HSF2 I Rep1, 12 for Halo-HSF2 I Rep2, 9 for Halo-HSF2 I Rep3, 11 for Halo-HSF2 M Rep1, 9 for Halo-HSF2 M Rep2, and 9 for Halo-HSF2 M Rep3. **(C)** H2B-corrected *k*_off_ values for HSF1-Halo and Halo-HSF2 constructs in interphase and mitosis, where each dot represents one biological replicate. Corrected *k*_off_ = *k*_off_(TF) - *k*_off_(H2B). Data depicted as mean ± SEM. P-values calculated using a standard two-tailed t-test with a 95% confidence interval.

**Supplemental Figure 5:**
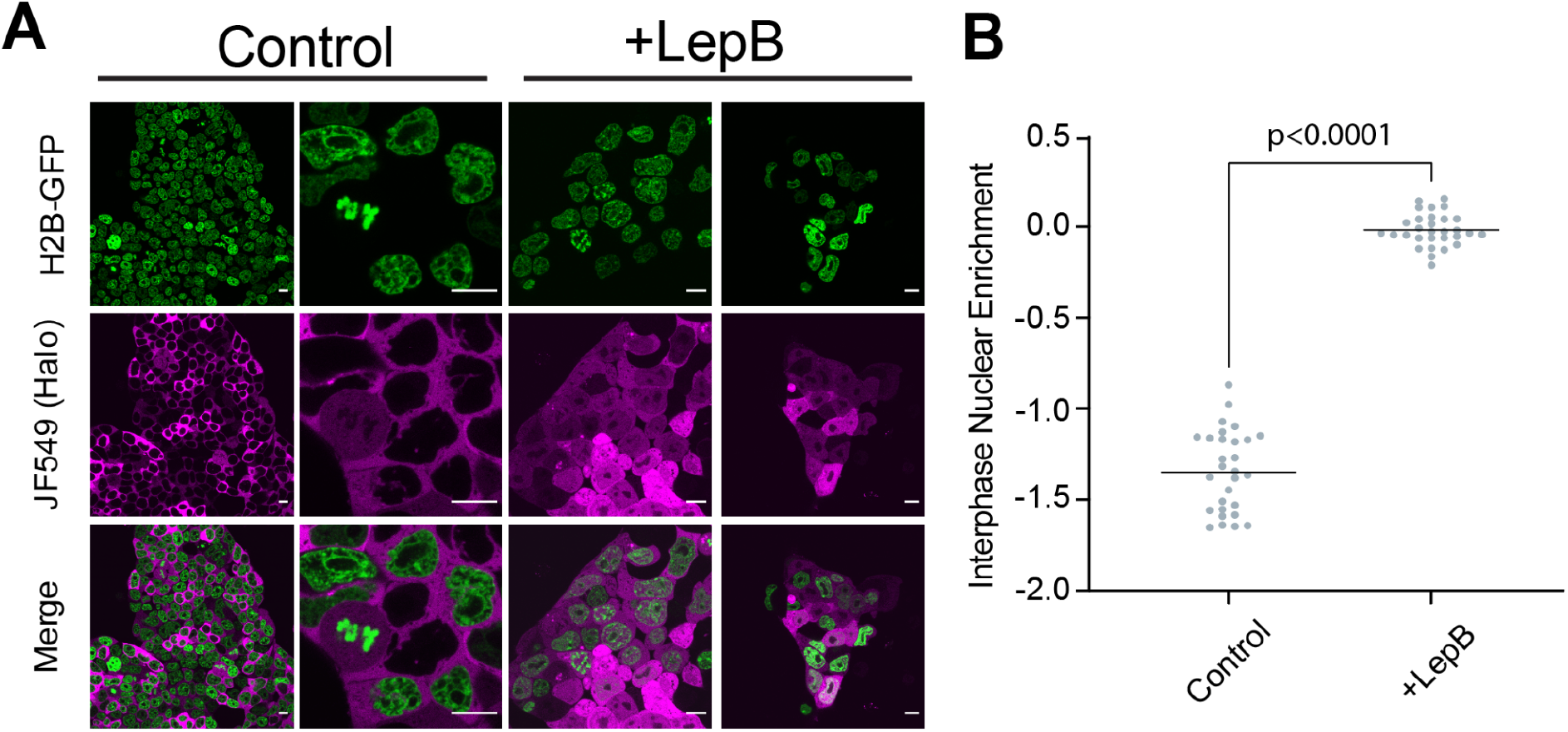
Validation of nuclear export inhibition. **(A)** Live-cell imaging of cells labeled with 200 nM JF549 dye (magenta) expressing Halo-NES construct. DNA is visualized by H2B-GFP overexpression (green). Scale bars are 10μm. Cells were either treated with 10 ng/mL leptomycin B (LepB) (+LepB) or with vehicle (Control) for 1h before imaging. **(B)** Nuclear enrichment quantification for the Halo-NES constructs (n=29 cells for NES-Halo control and 30 for NES-Halo +LepB across 3 biological replicates) in interphase cells. Data are visualized as individual data points with mean value indicated. P-values calculated using a standard two-tailed t-test with a 95% confidence interval.

**Supplemental Figure 6:**
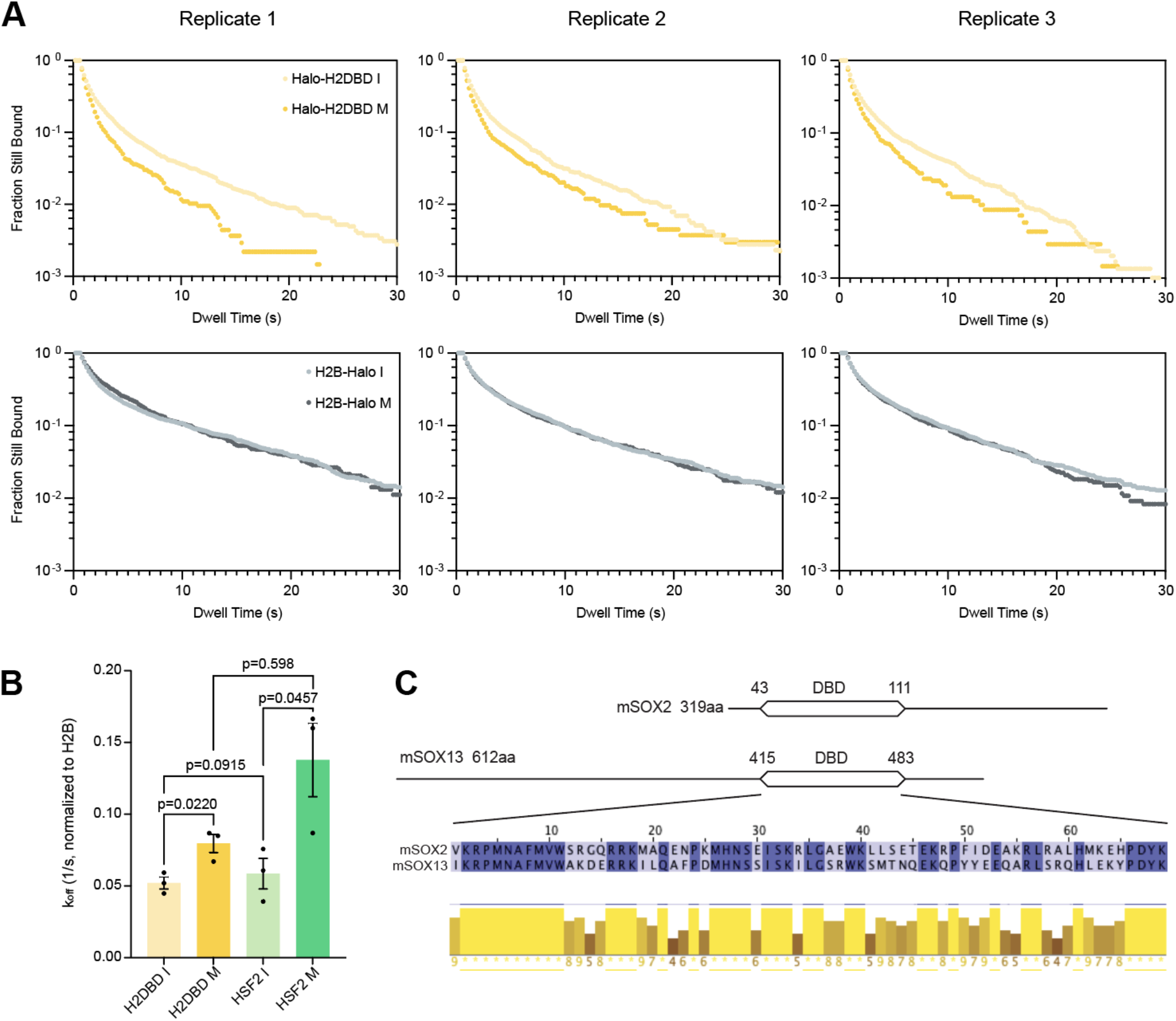
Replicate dwell curves for HSF1-Halo and Halo-HSF2 SPT, and SOX DBD sequence alignment. **(A)** Decay curves for each biological replicate of Halo-H2DBD (yellow, top) and the associated H2B-Halo replicate (gray, bottom) representing the fraction of HaloTagged molecules bound to DNA in interphase (I, lighter color) and mitosis (M, darker color) over time. Technical replicates: n=10 cells for H2B-Halo I Rep1, 10 for H2B-Halo I Rep2, 11 for H2B-Halo I Rep3, 10 for H2B-Halo M Rep1, 11 for H2B-Halo M Rep2, 12 for H2B-Halo M Rep3, 12 for Halo-H2DBD I Rep1, 11 for Halo-H2DBD I Rep2, 11 for Halo-H2DBD I Rep3, 11 for Halo-H2DBD M Rep1, 10 for Halo-H2DBD M Rep2, and 8 for Halo-H2DBD M Rep3. **(B)** H2B-corrected *k*_off_ values for Halo-HSF2 and Halo-H2DBD constructs in interphase and mitosis, where each dot represents one biological replicate. Corrected *k*_off_ = *k*_off_(TF) - *k*_off_(H2B). Data depicted as mean ± SEM. P-values calculated using a standard two-tailed t-test with a 95% confidence interval. **(C)** SOX2 DBD and SOX13 DBD sequence alignment (blue) and scoring matrices (yellow). Alignment produced with Jalview.

